# Sex differences in innate and adaptive neural oscillatory patterns predict resilience and susceptibility to chronic stress in rats

**DOI:** 10.1101/720011

**Authors:** Rachel-Karson Thériault, Joshua D. Manduca, Melissa L. Perreault

**Affiliations:** Department of Molecular and Cellular Biology, University of Guelph (ON)

## Abstract

Major Depressive Disorder (MDD) is a chronic illness with higher incidence in women. Dysregulated neural oscillatory activity is an emerging mechanism underlying MDD, however whether sex differences in these rhythms contribute to the development of MDD symptoms is unknown. Using the chronic unpredictable stress model, we found that stress-resilient and susceptible animals exhibited sex-specific oscillatory markers in the prefrontal cortex, cingulate cortex, nucleus accumbens and hippocampus. Resilient females were predominantly characterized by increased hippocampal theta power and coherence, while resilient males exhibited increased system-wide gamma coherence. In susceptible animals, the females displayed a widespread increase in delta and reduced theta power, however males showed few within-sex differences that could delineate stress susceptibility from resilience. Finally, stress responses were mediated by the temporal recruitment of specific neural pathways, culminating in system-wide changes that correlated with the expression of depression-like behaviours. These findings show that neurophysiological responses can serve as predictive markers of behaviours linked to depression in a sex-specific manner.

## Main

About 10% of the world population suffers from major depressive disorder (MDD), with the prevalence being two times greater in women than in men^1,2^. Chronic stress is an established risk factor for the development of MDD^3^, with clinical and preclinical studies showing females have greater behavioural sensitivity to chronic stress exposure^4–7^. This suggests that sex-differences exist within the stress response neural circuitry linked to the development of depression symptoms^8^. Despite the known predominance of depression in women, the majority of clinical and preclinical studies that have examined the influence of stress on depression have not employed sex as a factor. Thus, the mechanisms that contribute to increased female vulnerability to depression remain poorly understood.

The dysregulation of circuit function within the putative depression network has been linked to depression symptoms and antidepressant responsiveness, with many studies focusing on low frequency changes^9^. Electroencephalography (EEG) studies most commonly report frontal and parietal alpha asymmetries as potential endophenotypes of MDD^10–13^, as these patterns have been found to distinguish currently symptomatic and remitted patients from individuals with no history of depression^11, 12, 14, 15^. Alterations in midline theta activity in MDD are also commonly reported although results are not consistent, with some studies showing increased^16, 17^, decreased^18–21^ or no change^22,23^ in midline theta. A relationship between midline theta and antidepressant treatment response also exists but with the same discrepancies in findings^17, 20, 24–26^. Gamma band deficits are also implicated as a potential marker of MDD^27–29^ and therapeutic antidepressant efficacy^30–32^, with increased gamma associated with the remission of depression symptoms.

Few studies have considered the role of sex in dysregulated network activity in MDD, and of those that exist, there appears to be no consensus. For example, EEG studies have reported that the frontal alpha asymmetry is only present in women with MDD^33–36^ while others have shown it in both sexes^16, 37^. It has also been reported that only women have a positive association between frontal asymmetry and MDD severity^15, 33^. Similarly, whereas one study found that only women with MDD had increased right over left parietal activity^16^, another reported that women with MDD showed no differences in parietal activation compared to healthy controls^38^.

To better understand the role of neural oscillatory dysregulation in MDD, and the potential contribution to female stress vulnerability and depression, more information on how sex influences circuit function is required. We therefore evaluated sex differences in circuit function adaptations induced by chronic unpredictable stress (CUS) exposure in rats and discerned the ability of these functional signatures to serve as early markers of resilience or susceptibility. We found distinct sex-, regional-, and time-dependent differences in stress-induced neurophysiological patterns that exhibited low within group variability. Whereas the resilient and susceptible females exhibited distinct predictive patterns, the susceptible male rats were more clearly identified by a lack of expression of a male resilient signature. Overall, repeated stress exposure induced the temporal recruitment of circuits in susceptible and resilient animals, resulting in a system-wide network change that correlated directly to the expression of depression-like behaviours.

## Results

The CUS model of depression is considered to be the animal model of depression with the greatest validity and translational potential^39^. This is largely due to the fact that it can delineate sex differences as well as differences in stress vulnerability in rodents^6^. The present study was designed to acquire an approximate 50:50 cohort of stress-susceptible and -resilient animals within each sex. To distinguish between the two cohorts of animals, we employed two tests of depression-like behaviours (the forced swim test (FST) and the sucrose preference test (SPT) for despair and anhedonia, respectively). To be considered stress-susceptible, animals must have displayed significant changes from baseline in both the FST (60% increase in immobility) and the SPT (20% reduction in sucrose preference)^40^. At baseline, there were no sex differences in time spent immobile in the FST, while females showed lower sucrose preference, compared to males (Supplementary Fig. 1a, b). In the females, we observed that a maximum of 3 weeks of CUS exposure was sufficient to induce a depression-like phenotype in approximately half of the animals, whereas males required a maximum of 5 weeks of CUS exposure (Supplementary Fig. 2a, b). A test of anxiety-like behaviour (elevated plus maze (EPM)) was also included. At baseline, males exhibited greater anxiety-like behaviour, compared to females (Supplementary Fig. 1c), however following CUS exposure, no animals in either sex were found to have anxiety-like behaviour (Supplementary Fig. 2c). No behaviours were found to be correlated to the estrous cycle stage in female rats (Supplemental Fig. 3b, c). Further, cycling was significantly disrupted by week 2 of CUS in resilient and susceptible females (Supplemental Fig. 3n).

### Sex- and resiliency-dependent changes in stress-induced oscillatory power

Both clinical and preclinical evidence has demonstrated an important role for neural oscillations in depression^9^. Yet, to our knowledge, the literature that has evaluated sex differences in depression-associated oscillations is limited with inconsistent findings^41^. Therefore, we first wanted to characterize sex differences in baseline and stress-induced oscillatory power in regions implicated in depression, namely the prefrontal cortex (PFC), cingulate cortex (Cg), nucleus accumbens (NAc) and dorsal hippocampus (dHIP). At baseline we found no sex differences in spectral power within the PFC or Cg, however in the NAc, female rats showed lower beta power (Supplementary Fig. 1d-f). The majority of baseline sex differences were seen in the dHIP, as females displayed greater theta and beta power in this region, and lower delta and high gamma power, compared to males (Supplementary Fig. 1g). Further, the estrous cycle did not impact baseline spectral power in any brain region (Supplementary Fig. 3d-g).

We then examined the impact of CUS on oscillatory power and the associated sex- and resiliency-dependent differences from baseline measures. For all stress-susceptible animals, the LFP data from the day prior to their first expression of a depression-like phenotype was utilized for analysis. For all stress-resilient animals, data from the last day of CUS exposure was used. We observed prominent sex- and region-dependent changes in spectral power in response to stress exposure in susceptible and resilient animals (Fig. 1). Within the PFC, only susceptible females exhibited increased delta power (1-4 Hz) (P<0.0001) compared to baseline (Fig. 1a). Both susceptible males and females had reduced theta (6-10 Hz) power (M: P=0.04, F:P<0.0001), however this reduction was significantly larger in females (Fig. 1a). Further, the resilient females displayed increased beta (>12-30 Hz) (P>0.0001), low gamma (>30-60 Hz) (P=0.001) and high gamma power (>60-100 Hz) (P=0.001) following CUS exposure; changes not observed in males (Fig. 1b).

**Fig. 1:**
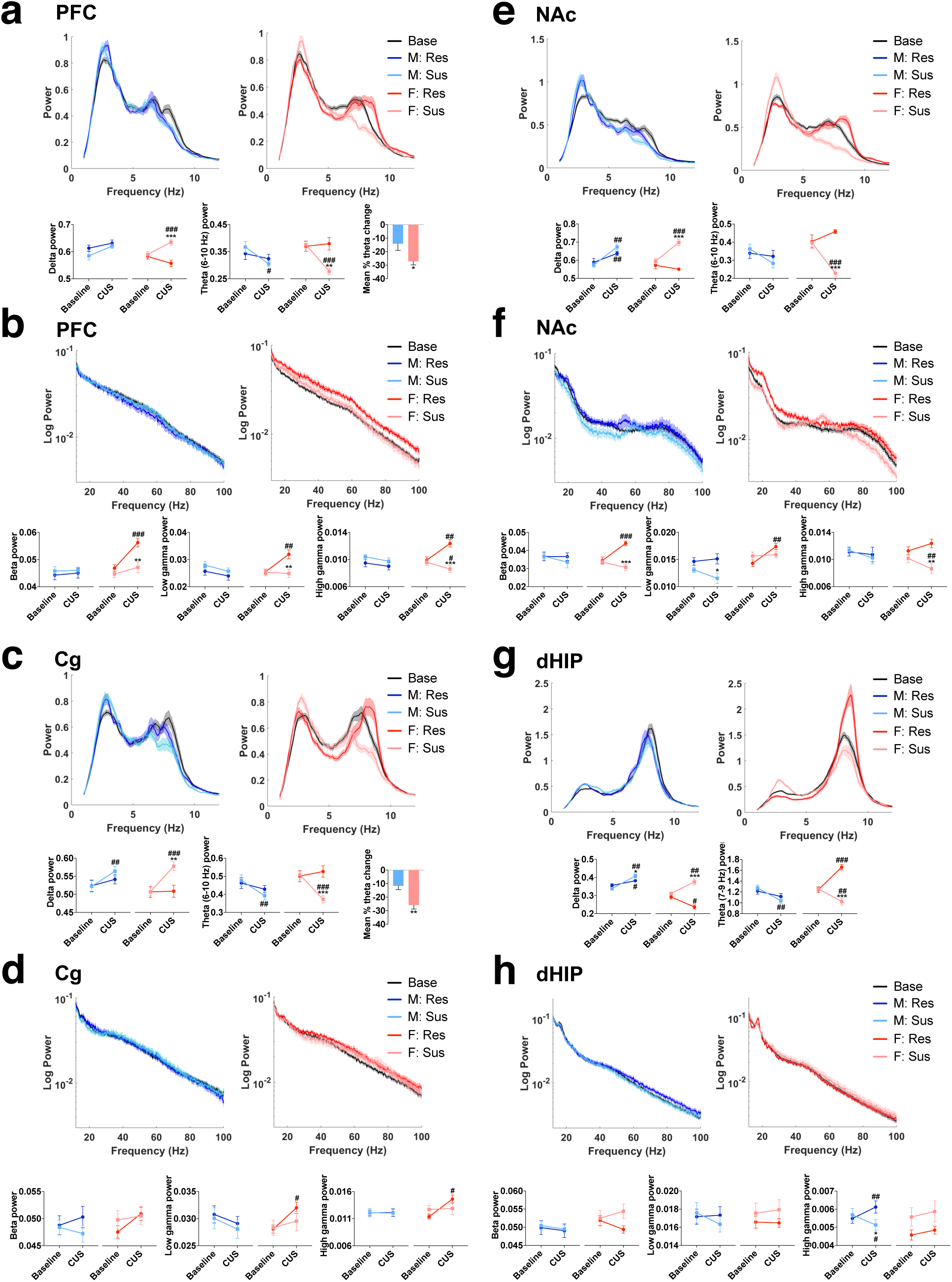
Sex- and region-dependent alterations in spectral power in response to CUS in susceptible and resilient rats. Normalized low and high frequency band power (top panels) and quantification of power spectra at each frequency (bottom panels). **a**, In the PFC, susceptible females exhibited increased delta power (Time x Resiliency x Sex interaction, *F*(1, 56)=7.977, P=0.007), while decreased theta (6-10 Hz) power was evident in both susceptible males and females (Time x Resiliency x Sex interaction, *F*(1, 58)=4.347, P=0.041). This theta reduction was more robust in female rats (*P<0.05 compared to male rats, Student’s *t* test, bottom right panel). **b**, Resilient females displayed increased power within all high frequency bands in the PFC (Time x Resiliency x Sex interaction, beta: *F*(1,59)=4.501, P=0.038; low gamma: *F*(1,61)=4.121, P=0.047; high gamma: *F*(1,55)=8.410, P=0.005). **c**, In the Cg, susceptible animals in both sexes exhibited increased delta (Time x Resiliency interaction, *F*(1, 60)=7.116, P=0.001) and reduced theta (6-10 Hz) power (Time x Resiliency x Sex interaction, *F*(1,60)=4.792, P=0.032). The decrease in theta power was more robust in female rats (**P<0.01 compared to male rats, Student’s *t* test, bottom right panel). **d**, Increased low and high gamma power was observed in resilient females (Time x Sex interaction, *F*(1,60)=8.053, P=0.005 and *F*(1,57)=9.583, P=0.003 for low and high gamma, respectively) in the Cg. **e**, In the NAc, susceptible females showed a reduction in theta (6-10 Hz) power (Time x Resiliency x Sex interaction, *F*(1,57)=11.176, P=0.001). **f**, Resilient females displayed an increase in beta (Time x Resiliency x Sex interaction, *F*(1,60)=4.039, P=0.049) and low gamma power (Time x Resiliency interaction, *F*(1,59)=8.93, P=0.004 and Time x Sex interaction, *F*(1,59)=4.775, P=0.033) in the NAc. **g**, In the dHIP, both susceptible males and females had reduced theta (7-9 Hz) power (Time x Resiliency x Sex interaction, *F*(1,53)=4.681, P=0.035), while theta power was increased in resilient females. **h**, High gamma power was increased and reduced in resilient and susceptible males, respectively in the dHIP (Time x Resiliency x Sex interaction, *F*(1,57)=9.035, P=0.004). Power spectra are presented as normalized data with jackknife estimates of SEM shown as shaded areas and with baseline data for each sex depicted by the black line. The quantified power spectra are presented as mean ± SEM. A three-way repeated measures ANOVA was used, unless otherwise indicated. *P<0.05, **P<0.01, ***P<0.001 compared to resilient, #P<0.05, ##P<0.01, ###P<0.001 compared to baseline. N=8-10/group, 2 electrodes/rat.

In the Cg, CUS induced enhanced delta power and reduced theta (6-10 Hz) power in both susceptible males (delta: P=0.004; theta: P=0.004) and females (delta: P<0.0001, theta: P<0.0001) (Fig. 1c). Similar to that observed in the PFC, this change in theta power was more robust in females, compared to males. Moreover, only resilient female rats exhibited increased low gamma (P=0.014) and high gamma power (P=0.001) from baseline following CUS exposure (Fig. 1d). Within the NAc, a stress-induced reduction in theta (6-10 Hz) power was observed in susceptible females only (P<0.0001) (Fig. 1e). Additionally, increased beta (P<0.0001) and low gamma power (P=0.006) was exclusively seen in resilient females (Fig. 1f). Lastly, in the dHIP, a reduction in theta power was once again observed in susceptible males (P=0.007) and females (P=0.004) following stress exposure although, unlike the other regions, this was restricted to the 7-9 Hz frequency band. Conversely, resilient females displayed increased theta (7-9 Hz) power (P<0.0001), in addition to reduced delta power (P=0.014) (Fig. 1g). Male-specific changes in high gamma power were also seen. Notably, a significant increase and reduction in high gamma power was found in resilient (P=0.004) and susceptible (P=0.036) males, respectively (Fig. 1h).

In summary, stress induced sex- and resiliency-dependent changes in oscillatory power in a region-specific manner. Overall, stress susceptibility in both sexes appeared to be characterized by low frequency power changes, with females showing more robust changes. However, stress resiliency was characterized by increased high frequency power in a sex-dependent manner (PFC, Cg, NAc in females and dHIP in males).

### Sex- and resiliency-dependent changes in stress-induced oscillatory coherence

As distinct resiliency- and sex-dependent stress-associated alterations in oscillatory power were observed, we next sought to evaluate stress-induced alterations in inter-regional communication through coherence (Fig. 2). Similar to that of spectral power, the stage of the estrous cycle was not found to influence baseline coherence (Supplementary Fig. 3h-m). At baseline, females had higher coherence in the delta, theta, and beta frequency bands in all region connections, except the Cg-NAc and PFC-dHIP, compared to males (Supplementary Fig. 1h-m). Further, at baseline, those females subsequently categorized as resilient exhibited higher theta (6-10 Hz) coherence within the dHIP connections, specifically in the PFC-dHIP (P=0.001), Cg-dHIP (P<0.0001) and NAc-dHIP (P=0.002), compared to females subsequently categorized as susceptible (Fig. 2a-c).

**Fig. 2:**
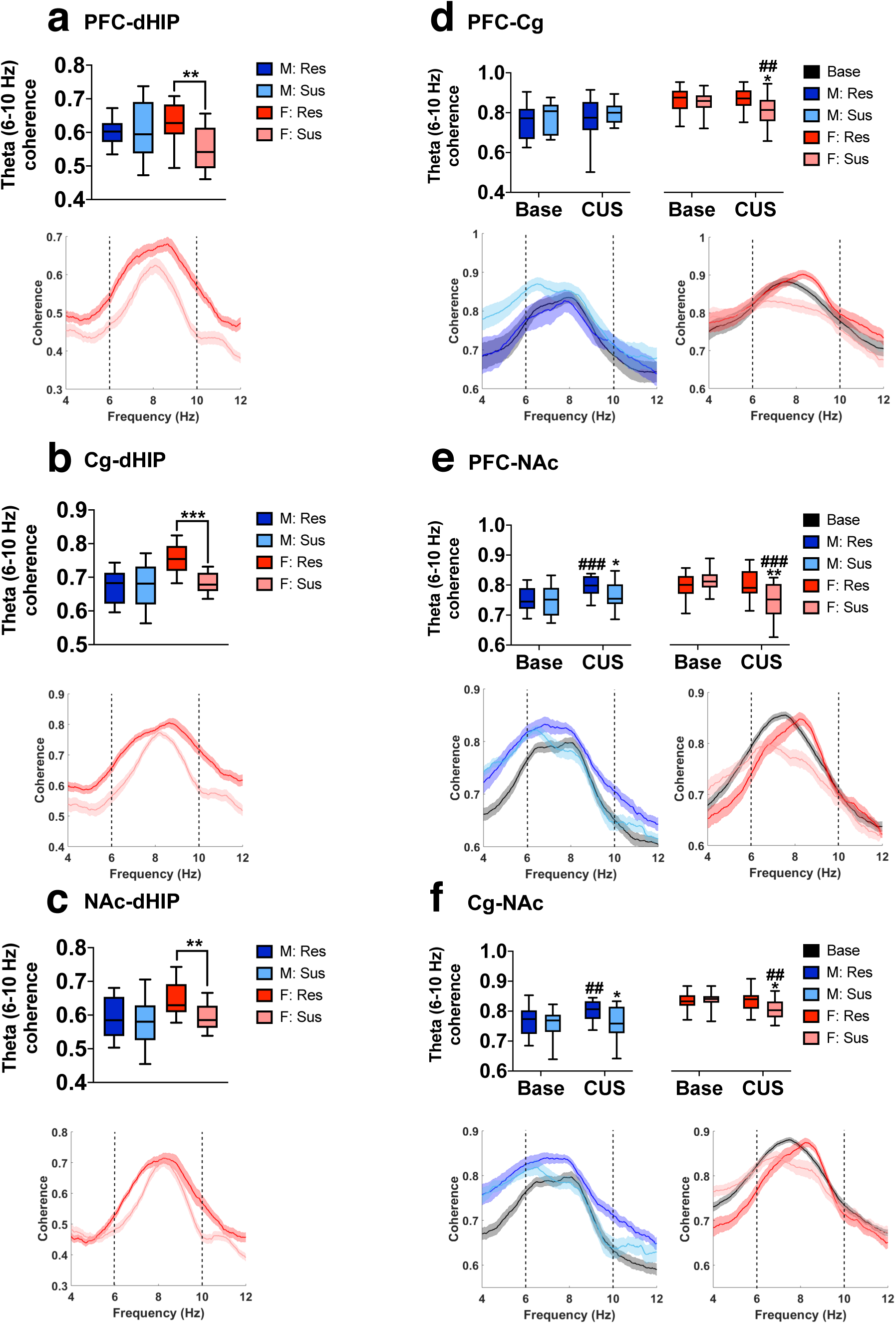
Sex-dependent alterations in theta frequency coherence at baseline and in response to CUS in susceptible and resilient rats. Sex-dependent CUS-induced changes in mean theta (6-10 Hz) frequency coherence (top panel) and theta coherence spectra (bottom panel). **a-c**, At baseline, resilient females exhibited greater theta (6-10 Hz) coherence between the PFC-dHIP (**a**), Cg-dHIP (**b**) and NAc-dHIP (**c**), compared to susceptible females (Student’s *t* test). **d**, In the PFC-Cg, susceptible females displayed reduced theta (6-10 Hz) coherence following CUS exposure (Time x Sex interaction, *F*(1,53)=7.094, P=0.01). **e, f**, Stress induced decreased and increased theta (6-10 Hz) coherence in susceptible females and resilient males, respectively, in the PFC-NAc (Time x Resiliency interaction, *F*(1,60)=27.482, P<0.0001 and Time x Sex interaction, *F*(1,60)=33.511, P<0.0001) (**e**) and Cg-NAc (Time x Resiliency interaction, *F*(1,57)=5.508, P=0.022 and Time x Sex interaction, *F*(1,57)=9.836, P=0.003) (**f**). Coherence spectra are presented as normalized data with jackknife estimates of SEM shown as shaded areas and with baseline data for each sex depicted by the black line, unless otherwise indicated. Quantified coherence is expressed as box-plots with min/max values shown. A three-way repeated measures ANOVA was used, unless otherwise indicated. *P<0.05, **P<0.01, ***P<0.001 compared to resilient, #P<0.05, ##P<0.01, ###P<0.001 compared to baseline. N=8-10/group, 2 electrodes/rat.

Following CUS exposure, theta (6-10 Hz) coherence alterations were observed in both sexes. In susceptible female rats, a stress induced a reduction in theta (6-10 Hz) coherence was observed between the PFC-NAc (P=0.002), PFC-Cg (P=0.028) and Cg-NAc (P=0.03). Conversely, resilient males exhibited a stress-associated increase in theta (6-10 Hz) coherence between the PFC-NAc (P=0.046) and Cg-NAc (P=0.038) (Fig. 2d-f). Resilient males also showed a distinct pattern of a global increase in high gamma coherence (PFC-NAc: P=0.028, PFC-Cg: P=0.003, PFC-dHIP: P<0.0001, Cg-NAc: P=0.004, Cg-dHIP: P=0.001, NAc-dHIP: P<0.0001) (Fig. 3). To summarize, results indicated that increased and reduced theta coherence is predominantly implicated in female stress resilience and vulnerability, respectively, while male resilience is more closely linked with a global upregulation in high gamma coherence.

**Fig. 3:**
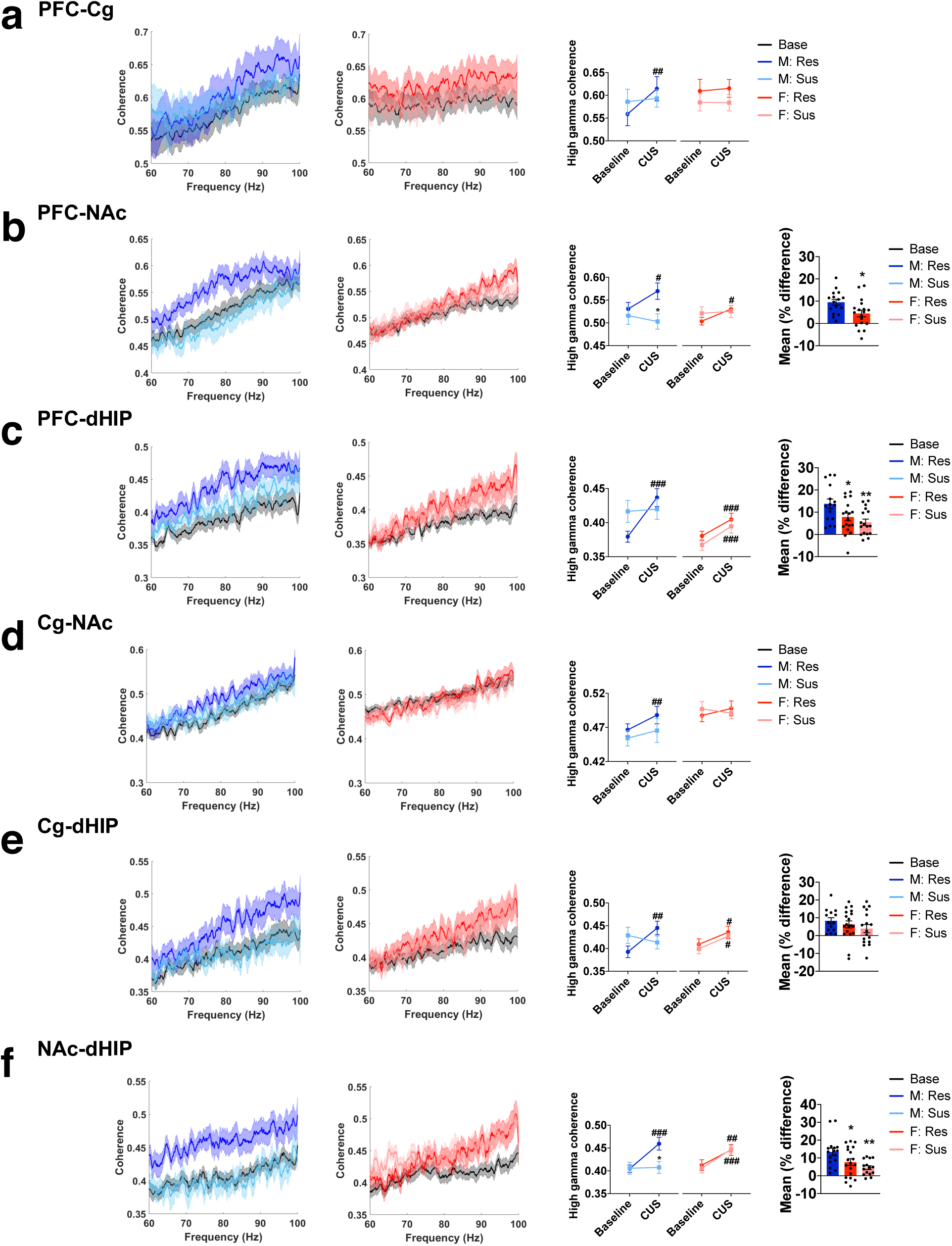
Sex-dependent alterations in high gamma frequency coherence following CUS exposure in susceptible and resilient rats. High gamma frequency coherence spectra for male (left panel) and female (centre panel) rats and mean coherence values (right panel). **a**, Between the PFC-NAc, only resilient males displayed increased high gamma coherence (Time x Resiliency interaction, *F*(1, 60)=7.673, P=0.007 and Sex x Resiliency interaction, *F*(1, 60)=3.816, P=0.05). **b**, Within the PFC-Cg, both resilient males and females exhibited increased high gamma coherence (Time x Resiliency interaction, *F*(1, 55)=5.771, P=0.02), however this change was more robust in male rats (*P<0.05 compared to male rats, Student’s *t* test, far right panel). **c**, Resilient males and females, as well as susceptible females had increased high gamma coherence between the PFC-dHIP (Time x Resiliency x Sex interaction, *F*(1, 56)=11.973, P=0.001) with greater effects observed in the male rats (*P<0.05 and **P<0.01 compared to male rats, Student’s *t* test, far right panel). **d**, Only resilient males exhibited increased high gamma coherence between the Cg-NAc (Time x Resiliency interaction, *F*(1, 53)=9.374, P=0.003 and main effect of Sex, *F*(1, 53)=8.448, P=0.005). **e, f**, Increased high gamma coherence was observed in resilient males and females, as well as susceptible females between the Cg-dHIP (Time x Resiliency x Sex interaction, *F*(1, 56)=4.729, P=0.034) (**e**) and NAc-dHIP (Time x Resiliency x Sex interaction, *F*(1, 57)=8.145, P=0.006) (**f**). Coherence spectra are presented as normalized data with jackknife estimates of SEM shown as shaded areas and with baseline data for each sex depicted by the black line. Quantified coherence is presented as mean ± SEM. A three-way repeated measures ANOVA was used, unless otherwise indicated. *P<0.05, **P<0.01, ***P<0.001 compared to resilient, #P<0.05, ##P<0.01, ###P<0.001 compared to baseline. N=8-10/group, 2 electrodes/rat.

### Sex-dependent correlations between stress-induced oscillatory changes and despair-like behaviour

Next, we wanted to determine whether the sex-specific stress-induced changes in oscillatory power and coherence observed were linked to the expression of a depression-like phenotype. Therefore, we correlated oscillatory changes with immobility time in the FST (Fig. 4 and 5), as it was the most robust behavioural change observed in susceptible animals following CUS. At baseline, female rats displayed a significant a negative correlation between theta (6-10 Hz) power and immobility in the PFC, Cg and NAc, and these relationships were strengthened following CUS exposure (Fig. 4a-c). However, no significant correlation was observed with delta or theta power in these regions in males (Supplemental Fig.4a-c). In the dHIP, however, delta and theta (7-9 Hz) power showed a significant positive and negative relationship with immobility, respectively, in both sexes, with a stronger relationship in females (Fig. 4d, e). Consistent with the reported stress-induced changes in dHIP high gamma power in males only, a male-specific negative correlation between high gamma power and immobility time was evident (Fig. 4f). Within the PFC and NAc, beta, low gamma and high gamma power were negatively correlated to immobility in females only (Supplemental Fig. 4d, f). Neither sex exhibited high frequency correlations in the Cg (Supplemental Fig. 4e).

**Fig. 4:**
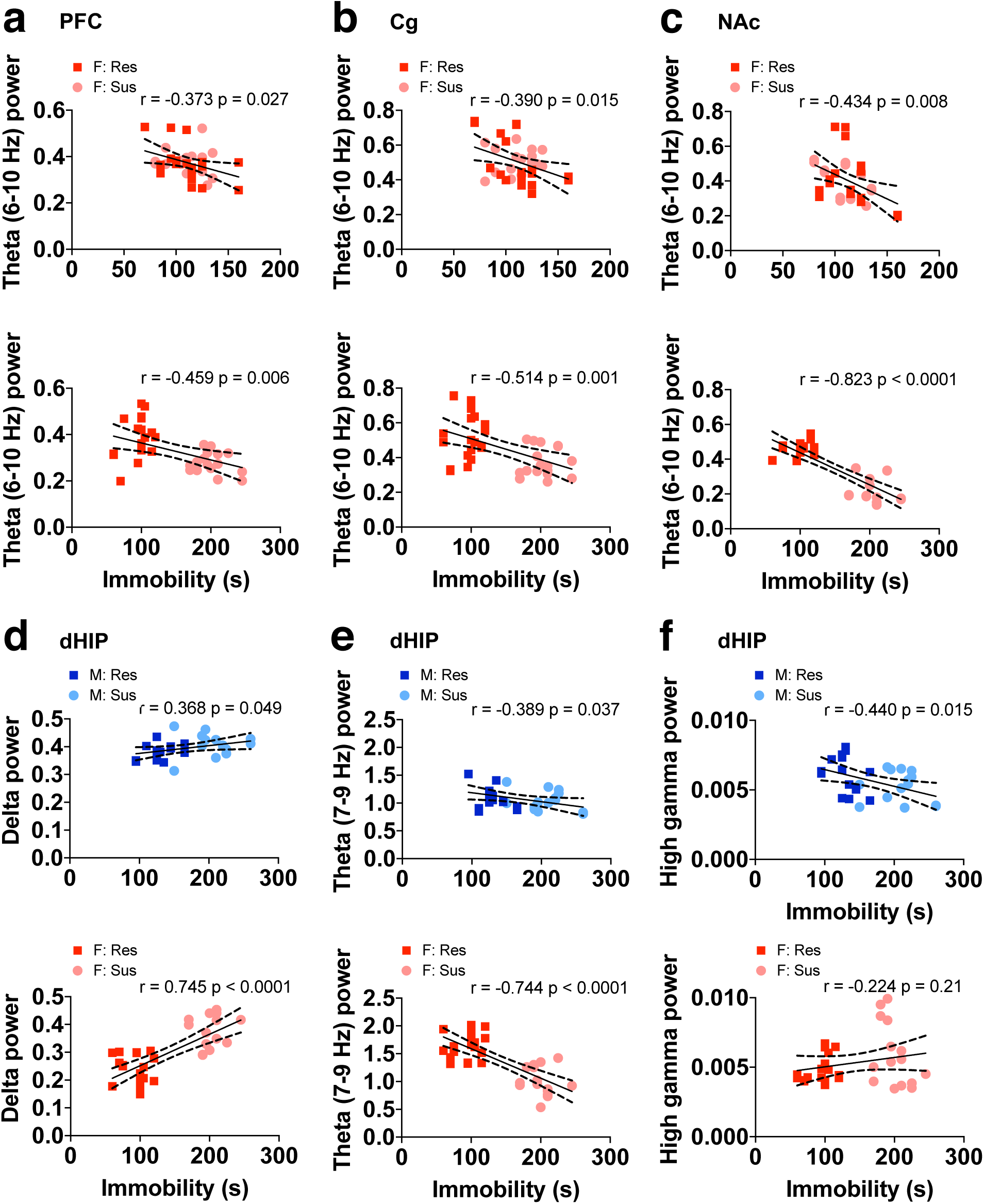
Sex-specific correlations between stress-induced alterations in spectral power and despair-like behaviour. **a-c**, Theta (6-10 Hz) power negatively correlated with time spent immobile in the forced swim test (FST) at baseline (top panel) and after CUS exposure (bottom panel) in the PFC (**a**), Cg (**b**) and NAc (**c**) of female rats only. **d, e**, In the dHIP, delta (**d**) and theta (7-9 Hz) (**e**) power positively and negatively correlated to immobility time, respectively, in both sexes, with a much stronger relationship observed in females. **f**, High gamma power in the dHIP correlated with immobility time in males only. All regression analyses were performed using Pearson’s correlation. All data is presented as individual data points with the line of best fit (solid line) ± SEM (dotted lines). N=8-10/group, 2 electrodes/rat.

**Fig. 5:**
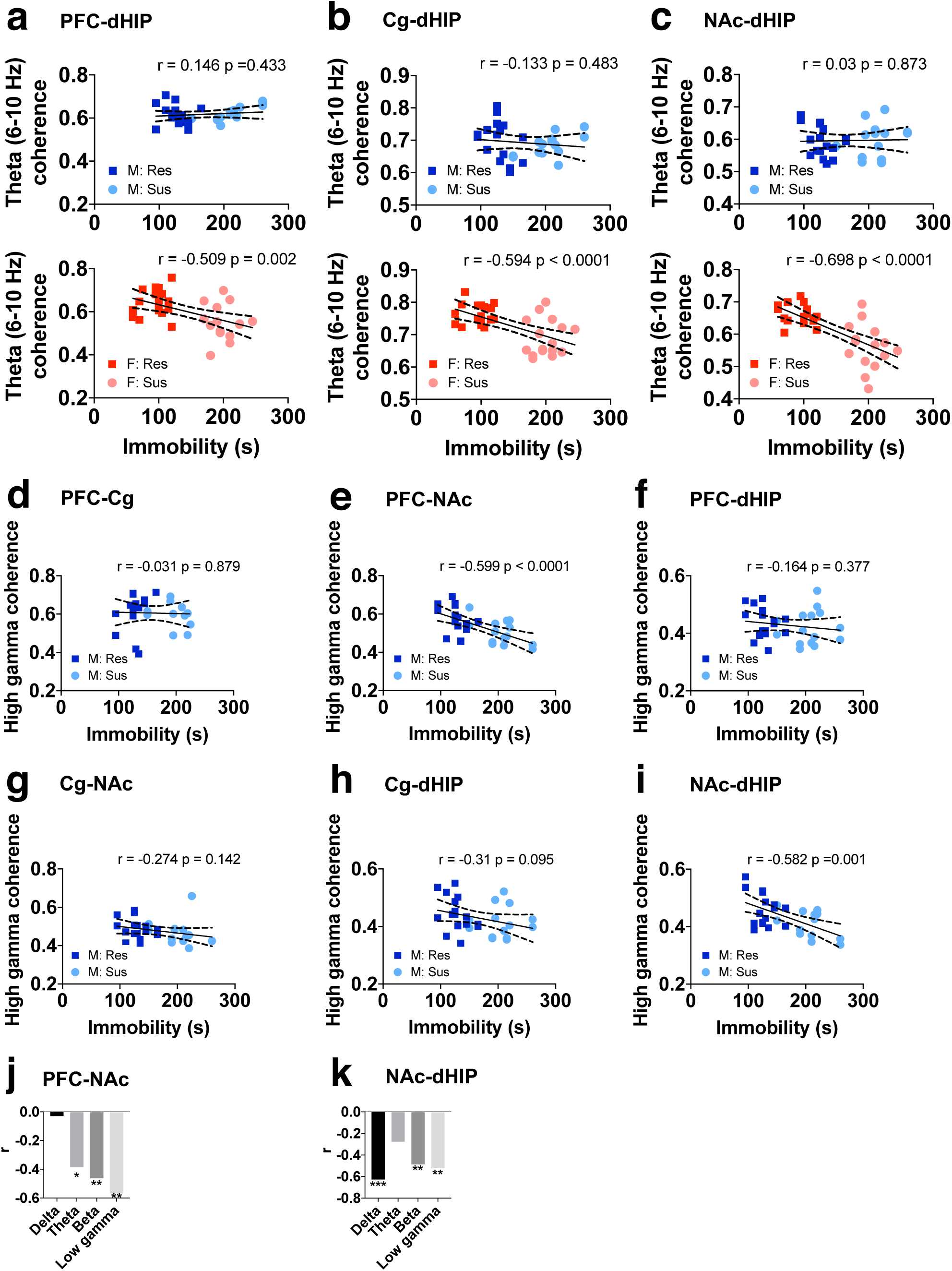
Sex-specific correlations between stress-induced alterations in coherence and despair-like behaviour. **a-c**, Theta (6-10 Hz) coherence negatively correlated with immobility time in the forced swim test (FST) in the PFC-dHIP (**a**), Cg-dHIP (**b**) and NAc-dHIP (**c**) in female rats only, following CUS exposure. **d-i**, High gamma coherence in males correlated with immobility time in the PFC-NAc (**e**) and NAc-dHIP (**i**), but not the PFC-Cg (**d**), PFC-dHIP (**f**), Cg-NAc (**g**) or Cg-dHIP (**h**) connections. **j**, Between the PFC-NAc, theta, beta and low gamma coherence in male rats negatively correlated with immobility time. **k**, Delta, beta and low gamma coherence between the NAc-dHIP of males negatively correlated with time spent immobile. All regression analyses were performed using Pearson’s correlation. **a-g**, data is presented as individual data points with the line of best fit (solid line) ± SEM (dotted lines). **j, k**, Data is presented as r value for each frequency band. *P<0.05, **P<0.01, ***P<0.001. N=8-10/group, 2 electrodes/rat.

Theta (6-10 Hz) coherence between the PFC-dHIP, Cg-dHIP and NAc-dHIP was negatively related to immobility in females, but not males (Fig. 5a-c). As high gamma coherence was globally increased in resilient males, we investigated whether this change was related to despair-like behaviour. Immobility negatively correlated to high gamma coherence between the PFC-NAc and NAc-dHIP of males (Fig. 5d-i) with no significant correlations in females (Supplemental Fig. 5a-f). Coherence between the PFC-NAc also correlated to immobility in the theta, beta and low gamma frequency bands and the NAc-dHIP coherence correlated in the delta, beta and low gamma frequency bands, in males only (Fig. 5j, k and Supplemental Fig. 5g, h). These findings indicate that the stress-induced alterations in spectral power and coherence were functionally related to the expression of behavioural despair in the FST following CUS exposure, in a sex-specific manner. In females, low frequency power in all regions and high frequency power in the PFC and NAc were most tightly coupled to behavioural despair, while in males, high gamma power in the dHIP and spectral coherence in the PFC-NAc and NAc-dHIP showed the strongest inverse relationship with behavioural despair following stress exposure.

### Predictive capacity of sex-dependent stress-induced oscillatory alterations

Our next goal was to delineate the time course during which these oscillatory changes occurred to determine whether early onset changes had the capacity to predict the subsequent development of a depression-like phenotype. As previously mentioned, LFP data collected 24 h prior to the first expression of a depression-like phenotype was used for stress-susceptible animals. Thus, for the purpose of the time course analyses, this day has been labeled as initial depression phenotype (IDP). All preceding time points are indicated by the number of days prior to IDP. For stress-resilient animals, IDP was considered the last day of CUS exposure.

In females, a pronounced distinction between stress-resilience and vulnerability was the observed changes in low frequency (delta and theta) power (Fig. 1). When investigated further, we found that theta (6-10 Hz) power alterations preceded those in the delta frequency band (Fig. 6a-d). Theta (6-10 Hz) power first decreased in the PFC of susceptible females, followed by the NAc at 9 days (P=0.01) and 7 days (P=0.035) prior to IDP, respectively (Fig. 6a, b). Reduced theta (6-10 Hz) power was last observed in the Cg (P=0.001) and the dHIP (P=0.004) at 4 days prior to IDP (Fig. 6c, d). Similarly, increased delta power in susceptible females was first seen in the PFC but occurred much later at 2 days prior to IDP (P=0.01) (Fig. 6a). This was followed by increased delta power occurring in the NAc (P<0.0001), Cg (P<0.0001) and dHIP (P=0.001) on IDP (Fig. 6b-d). In contrast, resilient females exhibited reduced delta power and increased theta (7-9 Hz) power in the dHIP (Fig. 1g) and these alterations were found to be an early adaptive stress response, occurring two days after the beginning of CUS exposure (i.e. IDP-16) (delta: P=0.015, theta: P<0.0001) (Fig. 6d). Low frequency power changes were observed in susceptible males as well, with significantly increased delta power and decreased theta (6-10 Hz) power observed 16 days prior to IDP in all brain regions (Supplemental Fig. 6a-d). However, these changes either did not persist or were also observed in resilient males.

**Fig. 6:**
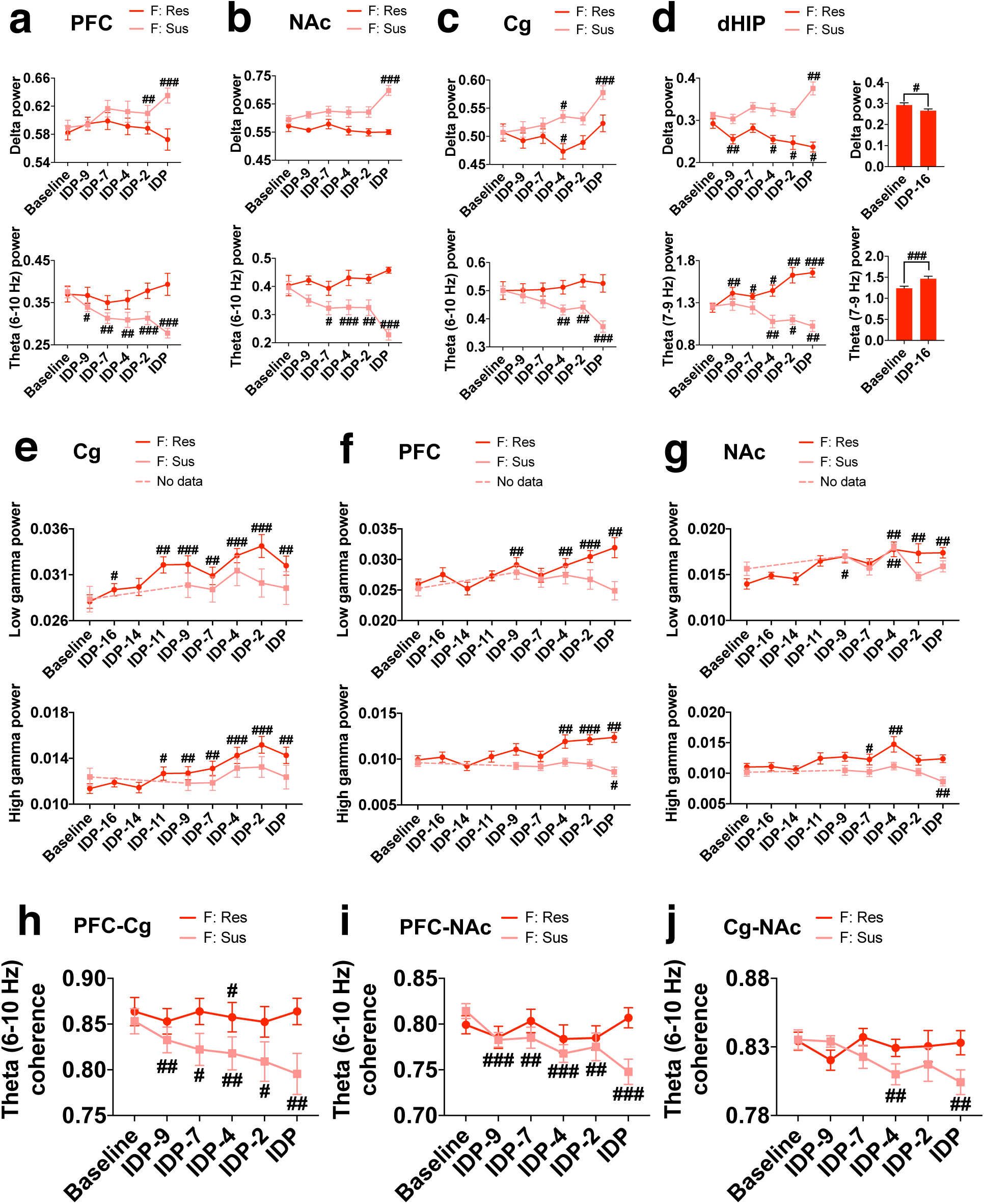
Time course of stress-induced oscillatory alterations in susceptible and resilient female rats. For stress-susceptible rats, LFP data collected 24 h prior to the first expression of a depression-like phenotype was used and has been labeled as initial depression phenotype (IDP). For stress-resilient animals, IDP was considered the last day of CUS exposure. All preceding time points are indicated by the number of days prior to IDP. **a-d**, In susceptible female rats, delta power (top panel) alterations proceeded those in the theta frequency (bottom panel). Both delta and theta power were first altered in the PFC (delta: Time x Resiliency interaction, *F*(5,125)=2.930, P=0.015; theta (6-10 Hz): Time x Resiliency interaction, *F*(5,110)=4.026, P=0.002) (**a**), then in the NAc (delta: Time x Resiliency interaction, *F*(5,135)=4.516, P=0.001; theta (6-10 Hz): Time x Resiliency interaction, *F*(5,140)=5.567, P<0.0001) (**b**) and lastly, in the Cg (delta: Time x Resiliency interaction, *F*(1,135)=2.513, P=0.033; theta (6-10 Hz): Time x Resiliency interaction, *F*(5,145)=4.668, P=0.001) (**c**). **d**, The dHIP was one of the last regions in which increased delta power and increased theta (6-10 Hz) power was observed in susceptible female rats (delta: Time x Resiliency interaction, *F*(5, 100)=3.428, P=0.007; theta (7-9 Hz): Time x Resiliency interaction, *F*(5,130)=7.333, P<0.0001), while resilient females exhibited opposite delta and theta (6-10 Hz) power changes after 48 h of stress exposure (on IDP-16). **e-g**, Stress-induced increases in low gamma (top panel) and high gamma (bottom panel) power in resilient females first occurred in the Cg (low gamma: main effect of Time, *F*(5, 160)=10.52, P<0.000; high gamma: Time x Resiliency interaction, *F*(5, 165)=2.654, P=0.02) (**e**), followed by the PFC (low gamma: Time x Resiliency interaction, *F*(5, 155)=4.255, P=0.001; high gamma: Time x Resiliency interaction, *F*(1, 145)=6.603, P<0.0001) (**f**). Low gamma alterations were last observed in the NAc (Time x Resiliency interaction, *F*(1, 120)=3.370, P=0.007) (**g**). **h-j**, Reduced theta (6-10 Hz) coherence in susceptible females occurred first between the PFC-Cg (Time x Resiliency interaction, *F*(5,145)=4.599, P=0.001)(**h**) and PFC-NAc (Time x Resiliency interaction, *F*(5, 145)=4.327, P=0.001) (**i**), followed by the Cg-NAc (**j**). All data is presented as mean ± SEM. A two-way repeated measures ANOVA was used, unless otherwise indicated. #P<0.05, ##P<0.01, ###P<0.001 compared to baseline. N=8-10/group, 2 electrodes/rat.

A key marker of female resilience appeared to be increased power in the gamma frequency bands within the PFC, Cg and NAc; changes not observed in males (Fig. 1). When the time course was examined, we found that both low and high gamma alterations first occurred in the Cg 16 (P=0.031) and 11 days (P=0.011) prior to IDP, respectively (Fig. 6e). Low gamma and high gamma power was next increased in the PFC of resilient females at 9 (P=0.001) and 4 days (P=0.007) prior to IDP, respectively (Fig. 6f). Low gamma power was also increased 9 days prior to IDP in the NAc (P=0.035) (Fig. 6g). Conversely, alterations in high gamma power within the dHIP were found to distinguish between stress resilient and susceptible males, with susceptible males exhibiting reduced high gamma power 16 days prior to IDP and resilient males showing increased high gamma power on IDP (Supplemental Fig. 6e).

Stress vulnerability in females was also marked by reduced theta (6-10 Hz) coherence between the PFC-NAc, PFC-Cg and Cg-NAc (Fig. 2d-f). When the timelines were examined, we observed that theta (6-10 Hz) coherence was first decreased between the PFC-NAc (P<0.0001) and the PFC-Cg (P=0.005) 9 days prior to IDP (Fig. 6h, i). This same reduction then occurred between the Cg-NAc 4 days prior to IDP (P=0.006) (Fig. 6j).

A prominent marker that accompanied male resilience to stress was a global increase in high gamma coherence (Fig. 3). We found that this alteration first occurred in resilient males in the NAc-dHIP 25 days prior to IDP (P=0.037) and was then observed in the PFC-dHIP, Cg-dHIP and PFC-Cg at 23 (P=0.042), 21 (P=0.04) and 18 (P=0.045) days prior to IDP, respectively (Fig. 7a-d). The Cg-NAc (P=0.01) and PFC-NAc (P=0.015) were the last connections in which greater high gamma coherence was seen in resilient males, both 2 days prior to IDP (Fig. 7e, f). Increased theta (6-10 Hz) coherence appeared to be another characteristic associated with male resiliency, specifically between the PFC-NAc and Cg-NAc (Fig. 2e, f). With further investigation, we found that theta (6-10 Hz) coherence first increased between the PFC-NAc 16 days prior to IDP (P=0.004) and then subsequently increased between the Cg-NAc on IDP (P=0.003) (Fig. 7g, h).

**Fig. 7:**
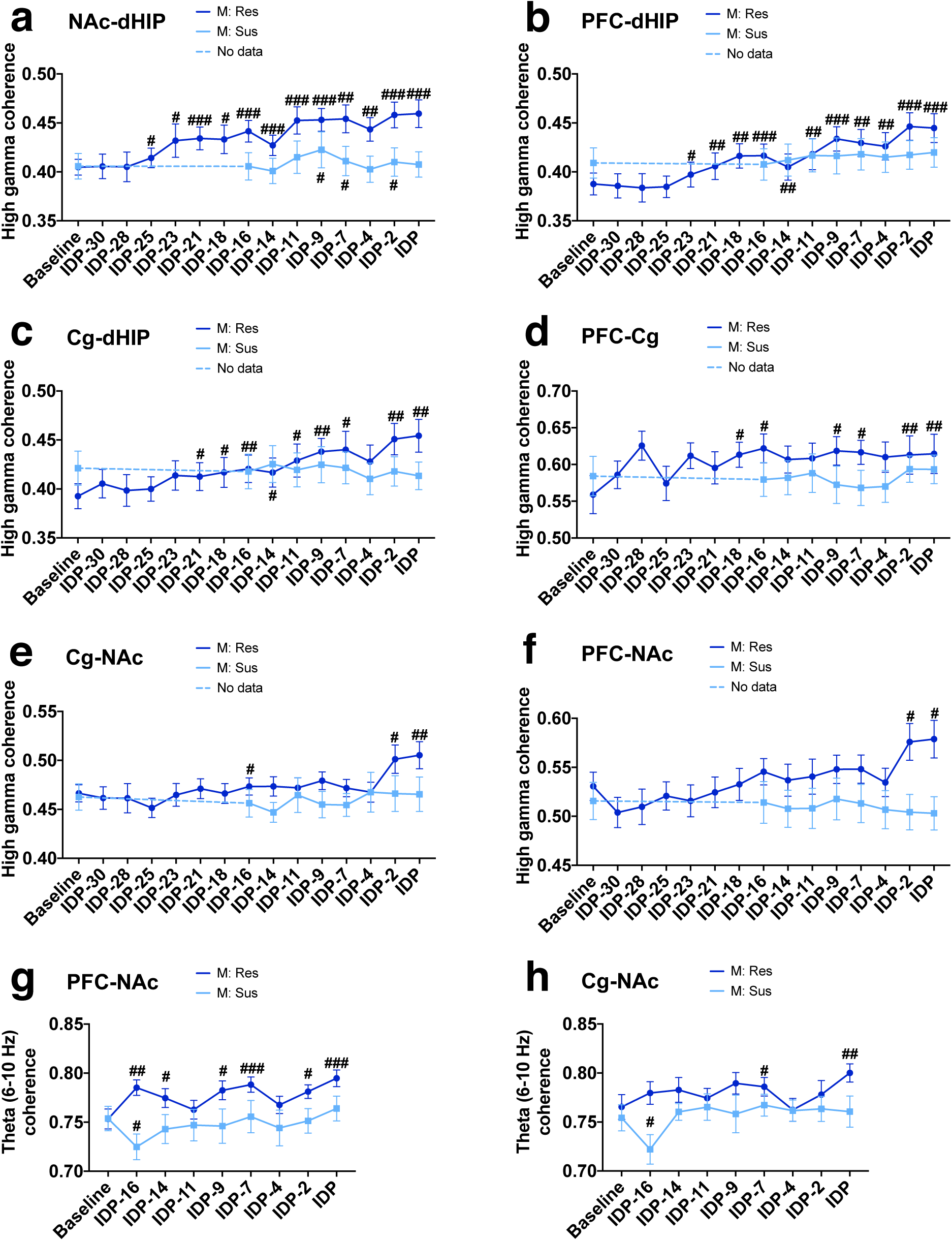
Time course of stress-induced oscillatory alterations in susceptible and resilient male rats. LFP data collected 24 h prior to the first expression of a depression-like phenotype was used for stress-susceptible rats and has been labeled as initial depression phenotype (IDP). All preceding time points are indicated by the number of days prior to IDP. IDP was considered the last day of CUS exposure for stress-resilient animals. **a-f**, In resilient males, stress-induced increased high gamma coherence first occurred between the NAc-dHIP (Time x Resiliency interaction, *F*(8, 176)=4.321, P<0.0001) (**a**), followed by PFC-dHIP (Time x Resiliency interaction, *F*(8,176)=3.564, P=0.001) (**b**), Cg-dHIP (Time x Resiliency interaction, *F*(8,184)=3.097, P=0.003) (**c**) and PFC-Cg (Time x Resiliency interaction, *F*(8,160)=1.934, P=0.05) (**d**). The last connections in which greater high gamma coherence was seen in resilient males were the Cg-NAc (Time x Resiliency interaction, *F*(8, 176)=2.176, P=0.031) (**e**) and PFC-NAc (Time x Resiliency interaction, *F*(8, 208)=2.417, P=0.016) (**f**). **g, h**, Resilient males were also characterized by increased theta (6-10 Hz) coherence following stress exposure, which first occurred between the PFC-NAc (Time x Resiliency interaction, *F*(8, 234)=2.458, P=0.014) (**g**) and Cg-NAc (main effect of Time, *F*(8,222)=2.218, P=0.027) (**h**). All data is presented as mean ± SEM. A two-way repeated measures ANOVA was used for all data. #P<0.05, ##P<0.01, ###P<0.001 compared to baseline. N=8-10/group, 2 electrodes/rat.

In summary, these results illustrated a sex-specific and time-dependent recruitment of brain regions and pathways associated with stress, that are indicated by specific changes in neural oscillatory systems function. Female resilient animals exhibited an innate elevation in coherence within the dHIP pathways (Fig. 2a-c) with early adaptive responses in low frequency (dHIP) and gamma (PFC, Cg, NAc) power (Fig. 6d-g). Conversely, male resilient animals showed early elevations in global high gamma coherence in response to stress (Fig. 7a-f). In susceptible animals, both sexes exhibited increased delta power and reduced theta power, although these changes were more robust and persistent in female animals (Fig. 6a-d and Supplemental Fig. 6a-d). For a full timeline of when sex- and resiliency-dependent oscillatory changes occurred, see Fig. 8.

**Fig. 8:**
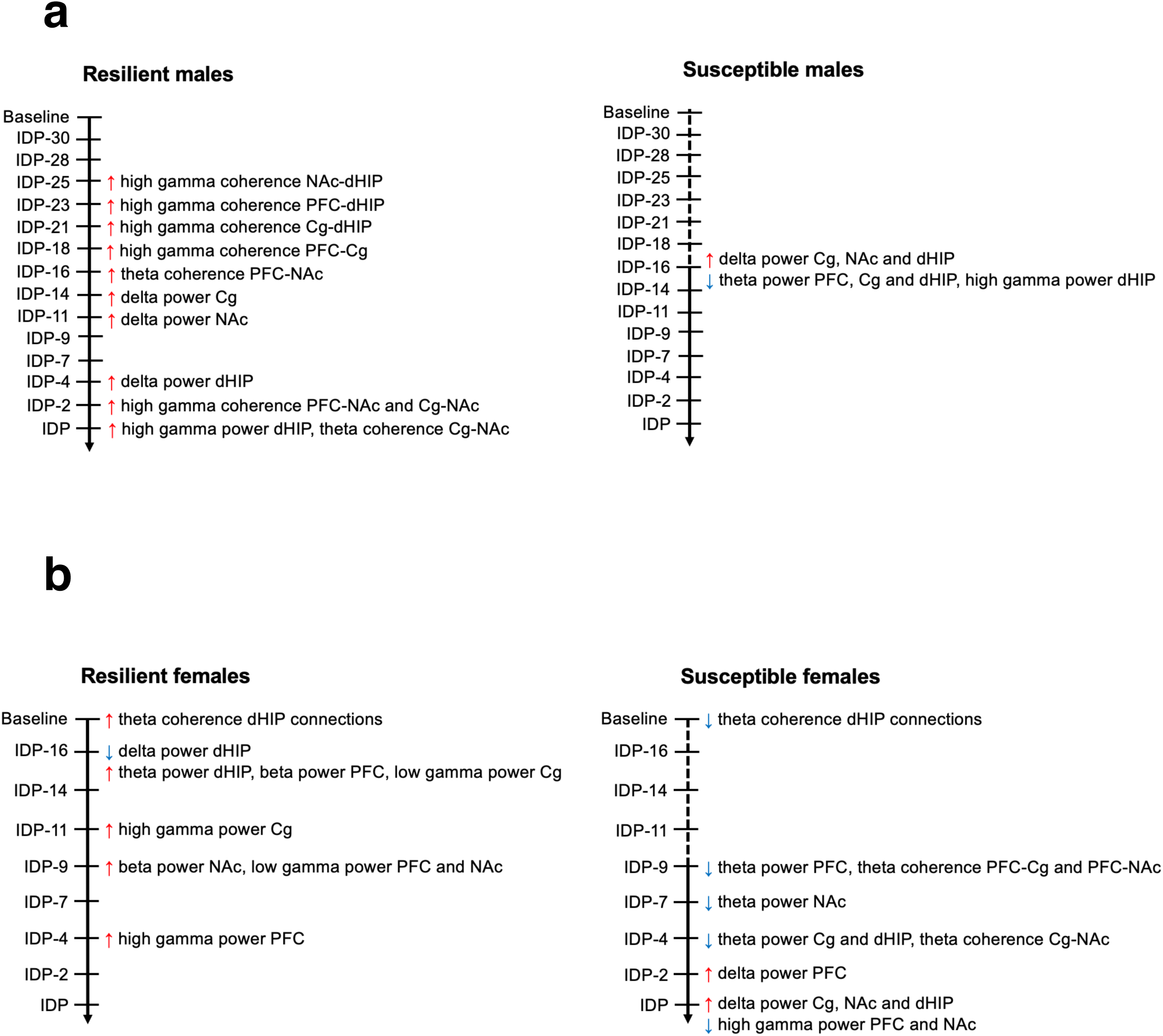
Time line of sex-dependent early oscillatory alterations predictive of resilience or susceptibility to chronic stress exposure in rats. Resilient (left panel) and susceptible (right panel) male and female rats exhibited distinct oscillatory alterations that preceded the manifestation (or absence) of depression-like behaviours. **a**, Resilient males were predominantly characterized by a global elevation in high gamma coherence, which was first observed between the NAc-dHIP after 7 days of stress exposure. This increased high gamma coherence then sequentially occurred in all other connections, suggesting that male resilience is dependent on the temporal recruitment of circuits that confers protection against chronic stress exposure. Conversely, susceptible males were largely marked by a lack of within-sex differences. Thus, it may be that markers of resilience are more critical in male stress responses, rather than markers of susceptibility. **b**, At baseline, resilient females could be distinguished from susceptible females due to their greater theta (6-10 Hz) coherence within dHIP connections. Resilient females also exhibited early low frequency power alterations in the dHIP, as well as increased high frequency power changes. In contrast, susceptible females displayed global increased and decreased delta and theta power, respectively, in addition to reduced theta (6-10 Hz) coherence between the PFC-Cg-NAc circuit. The timeline highlights the temporal dynamic of the adaptive changes in response to stress and that there are specific neurophysiological responses that confer resilience thereby preventing the onset of those oscillatory changes associated with susceptibility.

## Discussion

The present study aimed to elucidate sex differences in oscillatory patterns associated with mild chronic stress exposure in rats and the potential of these patterns to serve as early indices of susceptibility to develop depression-like behaviour. We demonstrated that male and female animals exhibited region-dependent differences in stress-induced neurophysiological patterns, with specific patterns predictive of susceptibility. Further, resiliency was not simply associated with a lack of susceptible signatures but manifested with distinct sex-specific innate and stress-induced oscillatory changes. Susceptibility or resiliency to repeated stress exposure was associated with a temporal recruitment of circuits, culminating in system-wide changes that reflected behavioural output.

Consistent with clinical^4, 5^ and preclinical^6, 7, 42^ reports, in the present study female rats were more susceptible to stress requiring a shorter duration of stress exposure to induce a depression-like phenotype compared to males. Stress-susceptible females could be distinguished from those that were resilient through differences in low frequency activity. Specifically, whereas susceptible females exhibited a system-wide increase in delta power that was concomitant with decreased theta power, the changes in the resilient females were restricted to the dHIP and opposite to that of the susceptible animals, with suppressed delta and increased theta power observed. A similar differentiation between susceptibility and resilience with respect to low frequency power was not apparent in the males, with all animals showing stress-induced increases and decreases in delta and theta, respectively, albeit with the effects more pronounced in the susceptible rats.

The interpretation of these findings is complex as sex differences in neural systems function in MDD have been poorly characterized^41^. Additionally, clinical studies that have examined low frequency activity in MDD report inconsistent findings, possibly reflective of the lack of sex as an inclusive experimental variable. In support of our findings, clinical EEG studies showed increased global or frontal delta activity in MDD patients^43–45^, as well as in women with menopausal depression^46^. With respect to theta oscillations, increased temporal theta and alpha power has been associated with improved mood^47^, whereas a global cortical reduction in theta and alpha power in MDD has been reported^48^. Reduced theta power in the occipito-parietal regions of MDD patients has also been hypothesized to reflect impaired affect regulation and functional connectivity^49, 50^ and indeed, cortical theta hypoconnectivity in MDD has been documented^45, 51^. In preclinical studies, a genetic mouse model of depression also showed reduced baseline theta oscillatory power in the prelimbic area of the PFC, which directly correlated with greater freezing time in the tail suspension test^28^. These findings demonstrating a link between reduced theta connectivity and MDD complement our own results showing pathway-specific and temporal increases in innate and stress-induced theta coherence in resilient female and male rats. It should be noted, however, that due to the chosen anatomical placements of our electrodes, important regional changes may not have been captured in the present study. For example, increased theta and alpha power has been observed in the subguenal Cg (infralimbic PFC of rats) and parietal cortex of depressed individuals^16^. However, reduced left midfrontal alpha hypoactivity, activity traditionally localized to the midfrontal Cg and more in line with our electrode placements, was also reported^16^ although theta/alpha asymmetries have yet to be documented in animal model systems of MDD. With consideration of these limitations, this study clearly demonstrated sex differences in CUS-induced low frequency oscillations that were temporally regulated in a region-dependent manner and which could delineate susceptibility from resilience. The notable sex differences in brain function in response to stress responsivity not only emphasizes that the manifestation of depression symptoms in males and females may be mediated by distinct mechanisms, but also critically highlights the need for inclusivity of both males and females in basic and clinical studies if we are to better understand the neuropathology of MDD.

Altered high frequency oscillatory activity was also linked to stress-susceptibility, and more strongly resilience. These changes were sex-specific, with susceptible females displaying reduced high gamma power in the PFC and NAc, resilient females displaying increased gamma frequency power in the PFC, Cg and NAc, and resilient males exhibiting a system-wide increase in high gamma coherence. A role for gamma oscillations in the neuropathology of MDD and antidepressant responsiveness has been repeatedly documented, although to our knowledge sex differences in these measures have not been previously evaluated. Individuals with MDD were shown to exhibit lower resting gamma activity in the rostral Cg compared to healthy individuals^27^, and previous preclinical findings are consistent with our own, showing reduced gamma power in the PFC and NAc being associated with a depression-like phenotype^28, 29^. Increased resting gamma power in MDD, particularly in the PFC, in response to repetitive transcranial magnetic stimulation has been linked to successful remission^30–32^, findings also supported by rodent studies showing that ketamine treatment to healthy rats increased gamma oscillatory activity in the PFC, NAc, amygdala, HIP and thalamus^52, 53^. Similarly, deep brain stimulation-induced normalization of suppressed high gamma coherence in the ventral tegmental area of Flinders rats, a model system of MDD, correlated with improved immobility measures in the FST^54^.

Our findings provide much needed insights into sex-specific, regional and temporal changes that occur in response to stress that emphasize distinct oscillatory markers linked to resilience and vulnerability. In females, we found that at baseline, a greater theta coherence across the dHIP connections could successfully distinguish between animals that were subsequently labeled as resilient and susceptible. Interestingly, 48 hours following stress exposure female stress resilience was accompanied by reduced delta power and increased theta power within the dHIP, with the opposite relationship observed in susceptible females. Notably, the aforementioned increases in gamma power displayed by the resilient females developed after the dHIP low frequency changes were established. Taken together, these findings indicate that the dHIP may function as a fulcrum for female stress resilience, where distinct low frequency alterations may be necessary to initiate events that trigger the observed increases in high frequency power within the PFC, Cg and NAc. In males, our findings suggest that it is the rapid and progressive increase in high gamma coherence in response to stress that contributed to resilience. Once again, resilience was associated with the temporal recruitment of all connections within the putative depression network, beginning first in the dHIP connections. Thus, although the neurophysiological mechanisms may differ, we hypothesize that it is the temporal recruitment of regions initiated by the dHIP that confers protection against stress exposure and the development of a depression-like phenotype.

Given the temporal dynamics of the observed adaptive changes associated with stress exposure, we posit that the neurophysiological responses that confer resilience play a key role in preventing the onset, or minimizing the functional impact, of those oscillatory changes associated with susceptibility, and that the nature of these changes are sex-specific. The importance of these resilience markers is perhaps the most obvious in the male animals where the resilient animals showed robust system-wide elevations in high gamma coherence, yet stable susceptibility markers in animals displaying a depression-like phenotype were not obvious. Specifically, although susceptible males showed stress-induced increases in delta and reductions in theta power in PFC, Cg and dHIP, the resilient rats followed a similar trend such that there appeared to be no differences between the two groups post-CUS. The only clear susceptibility marker we observed appeared to be a reduction in dHIP high gamma power, an effect opposite to that observed in the resilient males. Conversely, in the absence of resilience markers in female rats, the PFC appeared to be highly vulnerable to stress, being the first structure to exhibit stress-induced reductions in theta power in susceptible animals and these initial changes were soon followed by reduced theta power throughout the network. It is of note, however, that the appearance of many of these discrete susceptibility markers occurred several days prior to the initial expression of depression-like symptoms. This signifies that in females, oscillatory changes in discrete pathways may be insufficient to induce depression-like symptoms, with system-wide changes necessary for behavioural expression of the depression-like phenotype.

Finally, while no stress-induced reductions in NAc theta oscillations were observed in the male rats, female animals showed a significant coupling of baseline theta power in NAc with the immobility measure of behavioural despair, a relationship that strengthened significantly with stress exposure. Further, NAc low frequency power changes were most robust in susceptible females. The NAc has emerged as a critical player in MDD with key preclinical studies showing this region as being important in mediating stress responses in a sex-dependent manner. Subchronic stress exposure was reported to induce increased DNA methyltransferase 3 expression and enhanced activation of glutamatergic synapses in the NAc of female mice exhibiting a depression-like phenotype, while a more chronic stress protocol was required to induce the same alterations in male mice^55–57^. Thus, the NAc may mediate the female sensitivity to stress. Molecular markers of stress resilience have also been reported in the NAc^3, 58^. Recently, estrogen receptor α (ERα) was found to be a regulator of pro-resilient transcriptional changes in the NAc in both sexes, however males and females had minimal overlap in ERα downstream targets^58^. Further, an ovarian hormone-mediated upregulation of nuclear factor κB in the NAc was reportedly linked to stress-resilience, and therefore is a female-specific mechanism^3^.

In conclusion, our findings provide critically needed insights into sex-specific adaptive neurophysiological responses associated with stress and identify potential key oscillatory markers of resilience and susceptibility within regions of the depression network. In addition, stress exposure induces the sex-dependent temporal recruitment of circuits that result in the manifestation of depression-like behaviours in susceptible animals. As oscillatory markers of resiliency in both sexes occurred earlier with stress exposure than the changes observed in susceptible animals, we posit that the presence or absence of innate or stress-induced resilience-signatures is the most critical determinant of an animal’s subsequent stress response. Furthermore, our results highlight that stress influences neural oscillations in a sex-specific manner and as such, the inclusion of sex as an experimental factor and female animals in future clinical and preclinical studies, respectively, should be made a priority if we are to fully elucidate the neuropathology MDD and develop more effective treatments.

## Methods

### Animals

Adult male and female Wistar rats (Charles River, QC, Canada) weighing 175-225 g at the beginning of the experiment were used. Animals were housed in a temperature-controlled colony room, maintained on a twelve-hour reverse light/dark cycle (0700h lights off; 1900h lights on) with free access to water and food except during specific stages of stress exposure (See Chronic unpredictable stress). Behavioural testing was always conducted during the dark phase of the day-night cycle. Animals were handled daily for a minimum of 6 days prior to the start of the experiment. All procedures were approved by the Animal Care Committee of the University of Guelph and were carried out in accordance with the recommendations of the Canadian Council on Animal Care.

### Surgeries

Rats were anesthetized with isoflurane (5%), administered the analgesic carprofen (5 mg/kg, s.c.) and secured in a stereotaxic frame. Custom electrode microarrays were built using pre-fabricated Delrin templates and polyimide-insulated stainless steel wires (A-M Systems: 791600, 0.008”) and were implanted bilaterally into the medial region of the prefrontal cortex (PFC) (AP: +3.24 mm, ML: ±0.6 mm, DV: −3.8 mm), the anterior cingulate cortex (Cg) (AP: +1.9 mm, ML: ±0.5 mm, DV: −2.8 mm), the nucleus accumbens (NAc) (AP: +1.9 mm, ML: ±1.2 mm, DV: −6.6 mm) and the CA1 region of the dorsal hippocampus (dHIP) (AP: −3.5 mm, ML: ±2.5 mm, DV: −2.6 mm). A ground screw was implanted into the skull behind lambda and additional anchor screws were attached to the skull and secured with dental cement. All animals received a second carprofen injection 24 h following surgery and were allowed to recover in their home cage for a minimum of 7 days prior to experiments being performed. To verify electrode placements, lesions were produced using the Pulse Pal v2.0 (Sanworks) to deliver a 30 mv single train pulse through each electrode. Lesions were then visualized post-mortem (Supplementary Fig. 7).

### Electrophysiology

All LFP oscillatory recordings (Wireless 2100-system, Multichannel Systems) were performed in awake, freely moving animals in clear plexiglass boxes (18” x 18” x 18”). Baseline recordings were collected 24 h before the beginning of the CUS procedure and throughout the CUS protocol. Recordings were taken 3 times per week for 30 min and sampled at a rate of 1000 samples/s. Routines from the Chronux software package for MATLAB (MathWorks) were used to analyze the spectral power of LFP oscillations in each region and coherence between regions. Recordings were segmented, detrended, denoised and low-pass filtered to remove frequencies greater than 100 Hz. Continuous multitaper spectral power for the normalized data (to total spectral power) and coherence was calculated for delta (1-4 Hz), theta (> 4-12 Hz), beta (> 12-30 Hz), low gamma (> 30-60 Hz), and high gamma (> 60-100 Hz) unless otherwise stated.

### Apparatus and procedures

All female animals underwent vaginal lavage (100-200 μL 0.9% saline) to determine the stage of the estrous cycle prior to all behavioural testing and LFP recordings (Supplementary Fig. 3a). Baseline behaviours were collected prior to surgeries and behaviours were re-assessed weekly throughout the CUS procedure. All behavioural testing was conducted in a non-colony room isolated from external noise.

### Forced swim test

The forced swim test (FST) was used to assess behavioural despair. For the pre-test, animals were placed in a plexiglass cylinder with 24 ±1°C water filled to the height of 35 cm for 15 min. Animals were then towel-dried and placed back in their home cage. 24 h after the pre-test, animals were once again placed in the water-filled cylinder for 5 min. The water was changed after every animal. For all subsequent tests, the pre-test was not performed. At 5-s intervals, the following parameters were measured: climbing (both front paws breaking the surface of the water while attempting to jump out of the cylinder), swimming (movement of limbs paddling across the water surface) or immobility (passive floating with movements only necessary to keep nose above water).

### Elevated plus maze

Testing in the elevated plus maze (EPM) (Harvard Apparatus) was conducted to assess anxiety-like behaviours. The EPM was made of black plexiglass and was composed of a central square with two sets of opposing closed and open arms (50 cm x 10 cm). The closed arms were confined by 40 cm high walls along the edges, with the roof open. The entire maze was suspended 50 cm off the floor. Animals were placed in the centre square of the maze (head facing an open arm) and allowed to explore for 10 min. Behaviour was recorded and tracked using ANY-maze software (Stoelting Co.) and the following parameters were measured: the number of open and closed arm entries and total time spent in the open and closed arms.

### Sucrose preference test

The sucrose preference test (SPT) was used to assess anhedonia. Prior to the first test, two bottles of 1% (w/v) sucrose solution were placed in each cage for 24 h, in order for the animals to habituate to the sucrose solution. For the following 24 h, one bottle of sucrose solution was replaced with water. Once habituation was complete, animals were deprived of food and water for 24 h, after which the SPT was conducted. Animals were given two pre-weighed bottles: one containing 1% sucrose solution and one containing water. To minimise egocentric orientation bias, bottles were counterbalanced between cages and switched after every measurement. The two bottles were re-weighed every hour for 3 h. The percent sucrose preference (volume sucrose solution consumed/total volume consumed * 100) was calculated.

### Chronic unpredictable stress

All animals were exposed to chronic unpredictable stress (CUS) until half of the rats within each sex exhibited a depression-like phenotype. Animals that developed a depression-like phenotype were labeled as stress-susceptible and were characterized by a minimum 60% increase and 20% decrease from baseline in FST immobility and sucrose preference, respectively (Ho et al., 2017). Conversely, animals that did not show more than a 10% increase or 10% decrease in FST immobility and sucrose preference, respectively, from baseline were labeled as stress-resilient. The CUS protocol consisted of various uncontrollable physical, psychological and circadian stressors that were non-debilitating and inescapable. These stressors were: cold (4°C) exposure (1 h), intermittent on/off lights (8 h), cage tilt (30-45°) (8 h), damp bedding (500 mL water, 12-14 h), cold swim (10-13°C, 5 min), reverse light cycle (24 h), and food and water deprivation (24 h). Stressors were given on a random and unpredictable schedule and took place during both the light and dark cycle. The same stressor did not re-occur for at least 48 h and the stress schedules across each week were not identical. Throughout the CUS protocol, LFP recordings were collected 3 times per week and behavioural testing (FST, EPM, SPT) was re-assessed weekly (described above).

### Statistical analyses

Prior to all analyses, normality was assessed using the Shapiro-Wilk test. For all behavioural data, means were analysed using a repeated measures ANOVA with Resiliency as the between subjects factor and Time as the within subjects factor, followed by planned comparisons Student’s *t* test. For LFP data, routines from the Chronux software package (MATLAB, MathWorks) were used to extract individual frequency bands. Statistical analysis of LFP data was performed using a repeated measures ANOVA with Sex and Resiliency as between subjects factors and Time as the within subjects factor. In case of significant main effects or interactions, individual mean differences were identified by planned paired or independent Student’s *t* test to compare sex differences and stress-susceptibility effects at select time points, as within sex post-hoc comparisons are more representative of sex-specific stress impact. Analyses of the relationship between LFP data and behaviour were conducted using regression analyses with Pearson’s correlation. Computations were performed using IBM SPSS 24 software and are presented as the mean ± SEM.

## Acknowledgments

This research was supported by the Canadian Institutes of Health Research (grant number 450186 to MLP) and Vanier Canada Graduate Scholarship (RKT).

## Author contributions

MLP conceived the study and experimental design. RKT performed the behavioural experiments, stereotaxic surgeries and in vivo electrophysiological recordings with the help of JDM. RKT built the custom electrode arrays and conducted all MATLAB and statistical analyses. RKT and MLP wrote and edited the manuscript.

## References

1. McKenna, M.T., Michaud, C.M., Murray, C.J. & Marks, J.S. Assessing the burden of disease in the United States using disability-adjusted life years. Am J Prev Med 28,415–23 (2005).

2. Addis, M. E. Why is depression more prevalent in women? Clin Psychol Sci Pract 15, 153–168 (2008).

3. LaPlant, Q. et al. Role of nuclear factor κB in ovarian hormone-mediated stress hypersensitivity in female mice. Biol Psychiatry 65, 874–880 (2009).

4. Li, S. H. & Graham, B. M. Why are women so vulnerable to anxiety, trauma-related and stress-related disorders? The potential role of sex hormones. Lancet Psychiatry 4, 73–82 (2017).

5. Hodes, G. E. Sex, stress, and epigenetics: regulation of behaviour in animal models of mood disorders. Biol Sex Differ 4, 1 (2013).

6. Pfau, M. L. & Russo, S. J. Peripheral and central mechanisms of stress resilience. Neurobiol Stress 1, 66–79 (2015).

7. Rincon-Cortes, M. & Grace, A. A. Sex-dependent effects of stress on immobility behaviour and VTA dopamine neuron activity: Modulation by ketamine. Int J Neuropsychopharmacol 20, 823–832 (2017).

8. Shors, T. J., Chua, C. & Falduto, J. Sex differences and opposites effects of stress on dendritic spine density in the male versus female hippocampus. J Neurosci 21, 6292–9297 (2001).

9. Smart, O.L., Tiruvadi, V.R. & Mayberg, H. Multimodal approaches to define network oscillations in depression. Bio Psych 77,1061–1070 (2015).

10. Baskaran, A. et al. The comparative effectiveness of electroencephalographic indices in predicting response to escitalopram therapy in depression: A pilot study. J Affect Disord 227, 542–549 (2018).

11. Allen, J.J.B., Urry, H.L., Hitt, S.K. & Coan, J.A. The stability of resting frontal electroencephalographic asymmetry in depression. Psychophysiology 41, 269–280 (2004).

12. Henriques, J.B. & Davidson, R.J. Regional brain electrical asymmetries discriminate between previously depressed and healthy control subjects. J Abnorm Psychol 99, 22–31 (1990).

13. Domschke, K. et al. Magnetoencephalographic correlates of emotional processing in major depression before and after pharmacological treatment. Int J Neuropsychopharmacol 19, 1–9 (2016).

14. Gotlib, I.H., Ranganath, C. & Rosenfeld, J.P. Frontal EEG alpha asymmetry, depression, and cognitive functioning. Cogn Emot 12, 449–478 (1998).

15. Coan, J.A. & Allen, J.J.B. Frontal EEG asymmetry and the behavioral activation and inhibition systems. Psychophysiology 40, 106–114 (2003).

16. Jaworska, N., Blier, P., Fusee, W. & Knott, V. Alpha power, alpha asymmetry and anterior cingulate cortex activity in depressed males and females. J Psych Res 46, 1483–1491 (2012).

17. Korb, A.S., Hunter, A.M., Cook, I.A. & Leuchter, A.F. Rosteral anterior cingulated cortex theta current density and response to antidepressants and placebo in major depression. Clin Neurophysiol 120, 1313–1319 (2009).

18. Saletu, B., Anderer, P. & Saletu-Zyhlarz, G.M. EEG topography and tomography (LORETA) in diagnosis and pharmacotherapy of depression. Clin EEG Neurosci 41, 203–210 (2010).

19. Mitchell, D. J., McNaughton, N., Flanagan, D. & Kirk, I. J. Frontal-midline theta from the perspective of hippocampal “theta”. Prog Neurobiol 86, 156–185 (2008).

20. Mulert, C. et al. Rostral anterior cingulate cortex activity in the theta band predicts response to antidepressive medication. Clin EEG Neurosci 38, 78–81 (2007).

21. Mientus, S. et al. Cortical hypoactivation during resting EEG in schizophrenics but not in depressives and schizotypal subjects as revealed by low resolution electromagnetic tomography (LORETA). Psych Res 116 95–111 (2002).

22. Lubar, J.F., Congedo, M. & Askew, J. Low-resolution electromagnetic tomography (LORETA) of cerebral activity in chronic depressive disorder. Int J Psychophysiol 49, 175–185 (2003).

23. Pizzagalli, D. A. et al. Brain electrical tomography in depression: the important of symptom severity, anxiety, and melancholic features. Biol Psychiatry 15, 73–85 (2002).

24. Spronk, D., Arns, M., Barnett, K., Cooper, N. & Gordon, E. An investigation of EEG, genetic and cognitive markers of treatment response to antidepressant medication in patients with major depressive disorder: A pilot study. J Affect Disord 128, 41–48 (2011).

25. Arns, M. First EEG results of the iSPOT study in depression: EEG alpha asymmetry as a gender specific predictor of SSRI treatment outcome. Brain Stim 8, 337 (2015).

26. Broadway, J.M. et al. Frontal theta cordance predicts 6-month antidepressant response to subcallosal cingulate deep brain stimulation for treatment-resistant depression: a pilot study. Neuropsychopharmacology 37, 1764–1772 (2012).

27. Pizzagalli, D. A., Peccoralo, L. A., Davidson, R.J. & Cohen, J.D. Resting anterior cingulate activity and abnormal responses to errors in subjects with elevated depressive symptoms: a 128-channel EEG study. Hum Brain Mapp 27, 185–201 (2006).

28. Sauer, J.F., Struber, M. & Bartos, M. Impaired fast-spiking interneuron function in a genetic mouse model of depression. Elife 3, 4 (2015).

29. Voget, M. et al. Altered local field potential activity and serotonergic neurotransmission are further characteristics of the Flinders sensitive line rat model of depression. Behav Brain Res 15, 299–305 (2015).

30. Noda, Y. et al. Resting-state EEG gamma power and theta-gamma coupling enhancement following high-frequency left dorsolateral prefrontal rTMS in patients with depression. Clin Neurophysiol 128 424–432 (2017).

31. Bailey, N. W. et al. Responders to rTMS for depression show increased fronto-midline theta and theta connectivity compared to non-responders. Brain Stimul 11, 190–203 (2018).

32. Pathak, Y., Salami, O., Baillet, S., Li, Z. & Buston, C. R. Longitudinal changes in depressive circuitry in response to neuromodulation therapy. Front Neural Circuits 10, 50 (2016).

33. Stewart, J.L., Bismark, A.W., Towers, D.N., Coan, J.A. & Allen, J.J.B. Resting frontal EEG asymmetry an endophenotype for depression risk: sex-specific patterns of frontal brain asymmetry. J Abnorm Psychol 119, 502–512 (2010).

34. Bruder, G. E. et al. Electroencephalographic and perceptual asymmetry differences between responders and nonresponders to an SSRI antidepressant. Biol Psychiatry 49, 416–425 (2001).

35. Jesulola, E., Sharpley, C.F. & Agnew, L.L. The effects of gender and depression severity on the association between alpha asymmetry and depression across four brain regions. Behav Brain Res 321, 232–239 (2017).

36. Nusslock, R. et al. Comorbid anxiety moderates the relationship between depression history and prefrontal EEG asymmetry. Psychophysiology 55, e12953 (2018).

37. Jacobs, G.D. & Snyder, D. Frontal brain asymmetry predicts affective style in men. Behav Neurosci 110, 3e6 (1996).

38. Stewart, J.L., Coan, J.A., Towers, D.N. & Allen, J.J. Frontal EEG asymmetry during emotional challenge differentiates individuals with and without lifetime major depressive disorder. J Affect Disord 129 167–174 (2011).

39. Willner, P. The chronic mild stress (CMS) model of depression: History, evaluation and usage. Neurobiol Stress 6, 78–93 (2016).

40. Ho, Y. C. et al. Periaqueductal gray glutamatergic transmission governs chronic stress-induced depression. Neuropsychopharmacology 43, 302–312 (2018).

41. Theriault, R. K. & Perreault, M. L. Hormonal regulation of circuit function: sex, systems and depression. Biol Sex Differ 10, 12 (2019).

42. McFadden, L. M. et al. Sex-dependent effects of chronic unpredictable stress in the water maze. Physiol Behav 102, 266–275 (2011).

43. Bjork, M. H. et al. Quantitative EEG findings in patients with acute, brief depression combined with other fluctuating psychiatric symptoms: A controlled study from an acute psychiatric department. BMC Psychiatry 8, 89 (2008).

44. Morgan, M. L. et al. Influence of age, gender, health status, and depression on quantitative EEG. Neuropsychobiology 52, 71–76 (2005).

45. McVoy, M. et al. Resting-state quantitative electroencephalography demonstrates differential connectivity in adolescents with Major Depression Disorder. J Child Adolesc Psychopharmacol 29, 370–377 (2019).

46. Saletu, B. et al. Hormonal, syndromal and EEG mapping studies in menopausal syndrome patients with and without depression as compared with controls. Maturitas 23, 91–105 (1996).

47. Saletu, B. et al. Double-blind, placebo-controlled, hormonal, syndromal and EEG mapping studies with transdermal oestradiol therapy in menopausal depression. Psychopharmacology 122 321–329 (1995).

48. Shim, M., Im, C. H., Kim, Y. W. & Lee, S. H. Altered cortical functional network in major depression disorder: A resting-state electroencephalogram study. Neuroimage Clin 19, 1000–1007 (2018).

49. Flor-Henry, P., Lind, J.C. & Koles, Z.J. A source-imaging (low-resolution electromagnetic tomography) study of the EEGs from unmedicated males with depression. Psychiatry Res Neuroimaging 130, 191–207 (2004).

50. Pizzagalli, D.A., Oakes, T.R. & Davidson, R.J. Coupling of theta activity and glucose metabolism in the human rostral anterior cingulate cortex: An EEG/PET study of normal and depressed subjects. Psychophysiology 40, 939–949 (2003).

51. Iseger, T. A., Padberg, F., Kenemans, J.L., Gevirtz, R. & Arns, M. Neuro-cardiac-guided TMS (NCG-TMS): Probing DLPFC-sgACC-vague nerve connectivity using heart rate – First results. Brain Stimul 10, 1006–1008 (2017).

52. Hakami, T. et al. NMDA receptor hypofunction leads to generalized and persistent aberrant gamma oscillations independent of hyperlocomotion and the state of consciousness. PLoS One 4, e6755 (2009).

53. Hunt, M. J., Raynaud, B. & Garcia, R. Ketamine dose-dependently induces high-frequency oscillations in the nucleus accumbens in freely moving rats. Biol Psychiatry 60, 1206–1214 (2006).

54. Gazit, T. et al. Programmed deep brain stimulation synchronizes VTA gamma band field potential and alleviates depressive-like behaviour in rats. Neuropharmacology 91, 135–141 (2015).

55. Hodes, G. E. et al. Sex differences in nucleus accumbens transcriptome profiles associated with susceptibility versus resilience to subchronic variable stress. J Neurosci 35, 16362–16376 (2015).

56. Brancato, A. et al. Sub-chronic variable stress induces sex-specific effects on glutamatergic synapses in the nucleus accumbens. Neuroscience 350, 180–189 (2017).

57. Vialou, V. et al. Delta FosB in brain reward circuits mediates resilience to stress and antidepressant responses. Nat Neurosci 13, 745–752 (2010).

58. Lorsch, Z. S. et al. Estrogen receptor α drives pro-resilient transcription in mouse models of depression. Nat Commun 9, 1116 (2018).

