## Supplemental figures for "Sex differences in innate and adaptive neural oscillatory patterns predict resilience and susceptibility to chronic stress in rats"

**Supplemental Fig. 1: Baseline sex differences in depression-like behaviours and oscillatory activity in rats.**

**a-c,** No sex differences in time spent immobile in the FST at baseline were evident (**a**), however females showed a lower percent sucrose preference in the SPT (**b**) and spent more time in the open arms of the EPM (**c**). **d-g,** Power spectra (left and centre panels) and quantification of power spectra (right panel) at each frequency are shown. No sex differences in mean power spectra were observed in the PFC (**d**) or the Cg (**e**). **f,** In the NAc, female rats had lower beta power, compared to males. **g,** In the dHIP, female rats innately had lower delta and high gamma power, as well as greater theta and beta power, compared to male rats. **h-m,** Coherence spectra (left panel) and quantification (right panel) at each frequency are shown. **h,** Baseline coherence between the PFC-Cg of females was greater in the delta, theta, beta and low gamma frequency bands, compared to males. **i,** Within the PFC-NAc, coherence in the delta, theta and beta frequency bands was higher in female rats than male rats. **j,** In the PFC-dHIP connection, males innately had greater high gamma coherence. **k,** Baseline coherence of all frequency bands between the Cg-NAc were higher in females compared males. **l,** Within the Cg-dHIP, coherence in all frequency bands, except high gamma, were greater in females. **m,** Baseline theta frequency coherence was higher in females between the NAc-dHIP. Power and coherence spectra are presented as normalized data with jackknife estimates of SEM shown as shaded areas. Quantified power spectra and coherence are presented as percent difference from males ± SEM. Student's *t* test was used for all analyses. *P<0.05, **P<0.01, ***P<0.001, compared to males. N=17 males and N=19 females, 2 electrodes/rat.

**Supplemental Fig. 2: Behavioural validation of a CUS-induced depression-like phenotype.**

Animals that developed a minimum 60% increase and 20% decrease from baseline in FST immobility and sucrose preference, respectively, were labeled as stress-susceptible. Conversely, animals that did not show more than a 10% increase or 10% decrease in FST immobility and sucrose preference, respectively, from baseline were labeled as stress-resilient. **a,** In the FST, a significant increase in time spent immobile, indicative of despair-like behaviour, was observed in stress-susceptible males by week 4 of CUS (Time x Resiliency interaction, $F(5,75)=24.366$, $P<0.0001$), while the same effect was observed by week 2 of CUS in susceptible females (Time x Resiliency interaction, $F(3, 51)=33.903$, $P<0.0001$). **b,** In the SPT, susceptible male and female rats displayed reduced percent sucrose preference, indicative of anhedonia, by week 4 (Time x Resiliency interaction, $F(5, 75)=3.274$, $P=0.01$) and week 2 (Time x Resiliency interaction, $F(3, 51)=5.866$, $P=0.002$) of CUS, respectively. **c,** Neither susceptible or resilient males or females developed anxiety-like behaviour in the EPM due to CUS exposure. All data is presented as mean ± SEM and all analyses were completed using a two-way repeated measured ANOVA. *P<0.05, **P<0.01, ***P<0.001 compared to resilient. #P<0.05, ##P<0.01, ###P<0.001, compared to baseline. N= 8-10/group.

**Supplemental Fig. 3: Influence of estrous cycling on baseline behaviours and oscillatory activity in female rats.**

**a,** Representative photographs of cells collected during proestrus (left panel), estrus (left-centre panel), metestrus (right-centre panel) and diestrus (right panel) obtained using vaginal lavage. To ensure a sufficient sample size, rats in proestrus and diestrus were grouped together, as these stages are characterized by increased estradiol levels and rats in estrus and metestrus were grouped together as lower estradiol levels are present during these stages. **b, c,** Estrous cycling was not found to influence behavioural output in either the FST (**b**) or EPM (**c**). **d-g,** The stage of the estrous cycle did not impact baseline oscillatory power in any frequency band within the PFC (**d**), Cg (**e**), NAc (**f**) or dHIP (**g**). **h-m,** Baseline coherence in any frequency band was not affected by estrous cycling between the PFC-Cg (**h**), PFC-NAc (**i**), PFC-dHIP (**j**), Cg-NAc (**k**), Cg-dHIP (**l**) or NAc-dHIP (**m**). **n,** Length of estrous cycling significantly increased by week 2 of CUS exposure in resilient and susceptible female rats (#$P<0.05$, ##$P<0.01$, ###$P<0.001$ compared to baseline, paired t-test) **b, c, h-m,** Data is presented as mean ± SEM and **d-g,** data is represented as percent difference from diestrus/proestrus. All data was analyzed using a Student's *t* test, unless otherwise indicated.

**Supplemental Fig. 4: Sex- and region-dependent correlations between despair-like behaviour and CUS-induced alterations in oscillatory power.**

**a-c,** At baseline (top panel) and following CUS exposure (bottom panel) theta (6-10 Hz) power did not correlate with immobility time in the FST in the PFC (**a**), Cg (**b**) or NAc (**c**) of male rats. **d-f,** Correlations between time spent immobile and beta power (left panel), low gamma power (centre panel) and high gamma power (right panel) in male and female rats are shown. **d,** In the PFC, females, but not males, exhibited a relationship between oscillatory power at all high frequency bands and despair-like behaviour in the FST. **e,** In the Cg, beta, low gamma and high gamma power did not correlate with immobility time in either sex. **f,** Within the NAc, beta and high gamma power correlated with time spent immobile in females only, whereas low gamma power correlated with immobility time in both sexes. Data is presented as individual data points with the line of best fit (solid line) ± SEM (dotted lines). All regression data was analyzed using Pearson's correlation. N=8-10/group, 2 electrodes/rat.

**Supplemental Fig. 5: Region-dependent correlations between despair-like behaviour and coherence in female rats.**

**a-f,** No significant relationship between spent immobile in the FST and high gamma coherence in the PFC-Cg (**a**), PFC-NAc (**b**), PFC-dHIP (**c**), Cg-NAc (**d**), Cg-dHIP (**e**) or NAc-dHIP (**f**) connections was observed in female rats. **g,** Between the PFC-NAc of females, only coherence in the delta frequency band was correlated to immobility time. **h,** In the NAc-dHIP connection of females, time spent immobile only correlated to theta coherence in female rats. **a-f,** Data is presented as individual data points with the line of best fit (solid line) ± SEM (dotted lines). **g, h,** Data is presented as r value for each frequency band. All regression data was analyzed using Pearson's correlation. N= 9-10/group, 2 electrodes/rat.

**Supplemental Fig. 6: Time course of stress-induced alterations in oscillatory power in susceptible and resilient rats.**

Initial depression phenotype (IDP) represents LFP data collected 24 h prior to the first expression of a depression-like phenotype in stress-susceptible rats. For stress-resilient animals, IDP was considered the last day of CUS exposure. All preceding time points are indicated by the number of days prior to IDP. **a,** In the PFC, susceptible males exhibited an increase in delta power (Time x Resiliency interaction, $F(8, 160)=2.488$, $P=0.014$) and decrease in theta (6-10 Hz) power (main effect of Time, $F(8, 152)=3.304$, $P=0.002$) 16 days prior to IDP, however the altered delta power did not persist until the end of CUS exposure. **b,** In the Cg, increased delta power (Time x Resiliency interaction, $F(8, 182)=1.990$, $P=0.05$) and decreased theta (6-10 Hz) power (main effect of Time, $F(8, 160)=4.530$, $P<0.0001$) were observed 16 days prior to IDP in susceptible males, but not resilient males. **c,** In the NAc, stress-induced an early and persistent increase in delta power in both resilient and susceptible males (Time x Resiliency interaction, $F(8, 176)=3.178$, P=0.002), whereas stress-induced an early decrease in theta (6-10 Hz) power in both resilient and susceptible males (Time x Resiliency interaction, $F(8, 168)=2.102$, P=0.038), but this effect did not persist until the end of CUS exposure. **d,** In the dHIP, delta power was increased in both resilient and susceptible males (main effect of Time, $F(8, 160)=4.671$, P<0.0001), however this alteration occurred earlier in susceptible males. On the other hand, theta (7-9 Hz) power showed an early and persistent reduction in susceptible males only (main effect of Time, $F(8, 152)=3.5881$, P=0.001). **e,** High gamma power in the dHIP increased in resilient males on IDP (Time x Resiliency interaction, $F(8, 192)=2.492$, P=0.014), while no alterations in dHIP high gamma power were observed in females. All data is presented as mean ± SEM. A two-way repeated measures ANOVA was used for all data. #P<0.05, ##P<0.01, ###P<0.001 compared to baseline. N=8-10/group, 2 electrodes/rat.

**Supplemental Fig. 7: Histological verification of electrode placement.**

**a-d,** Electrode verification in the PFC (**a**), Cg (red arrow) (**b**), NAc (**c**) and dHIP (**d**).

### Supplemental Fig. 1

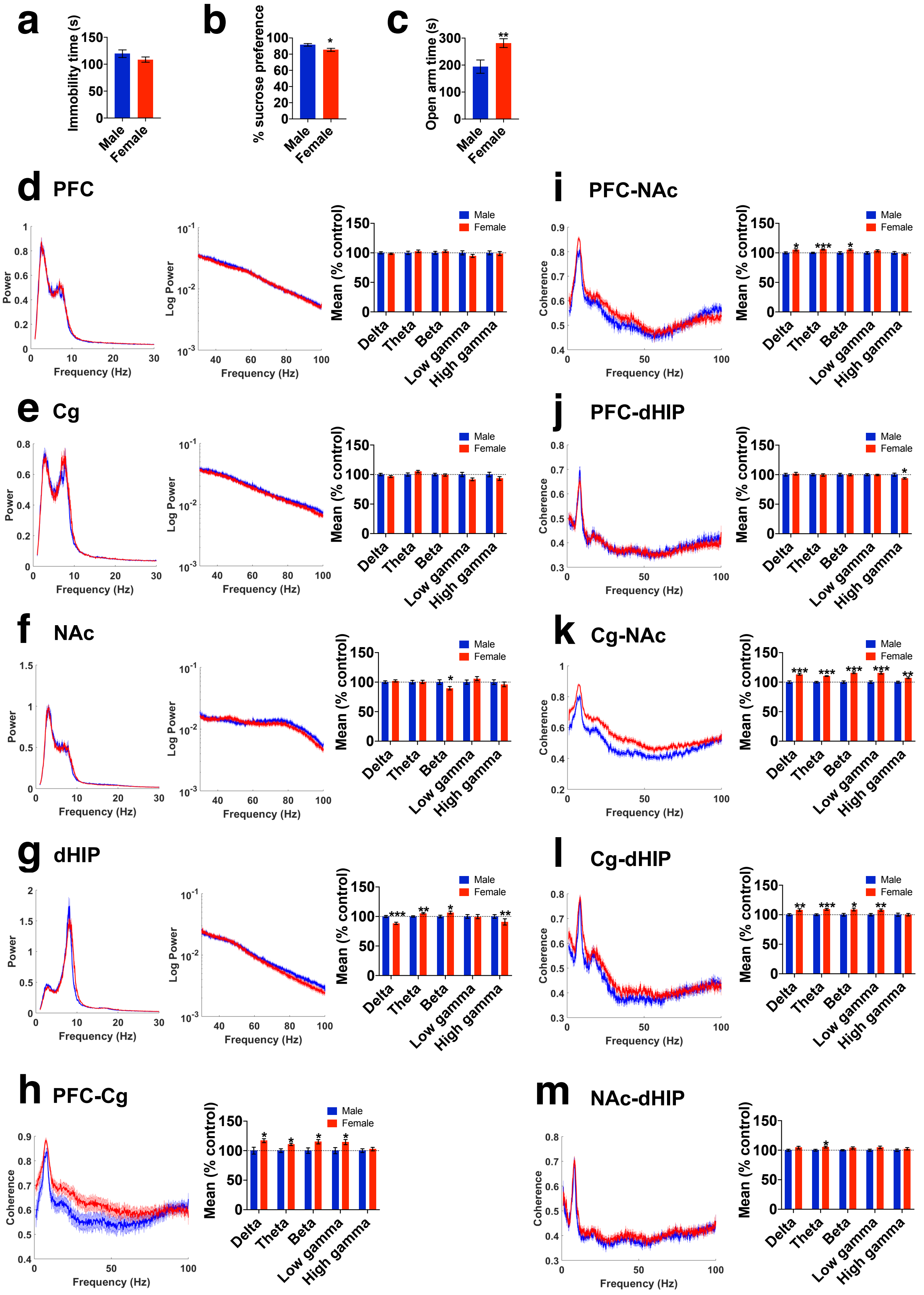

Supplementary Fig. 2

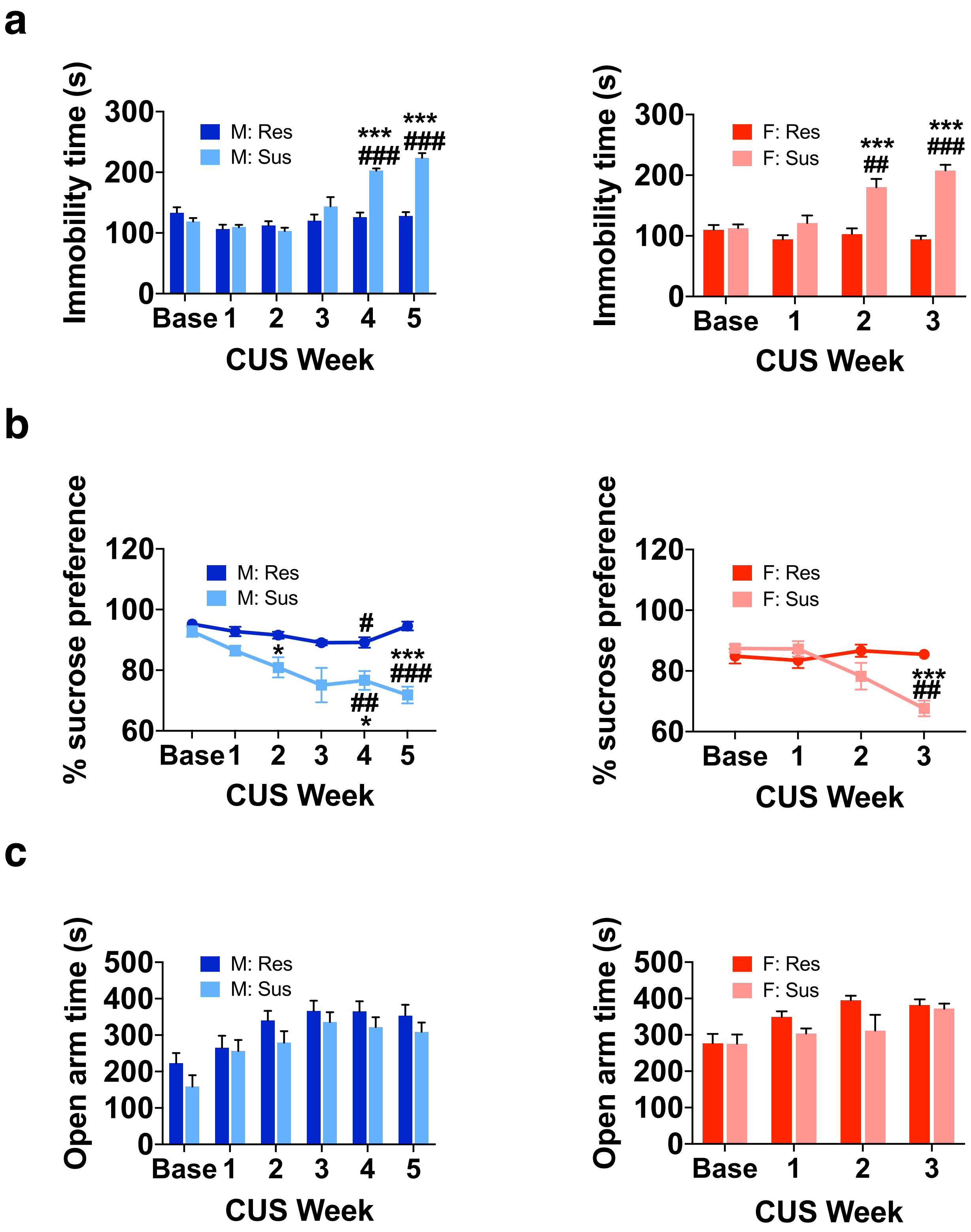

Supplemental Fig. 3

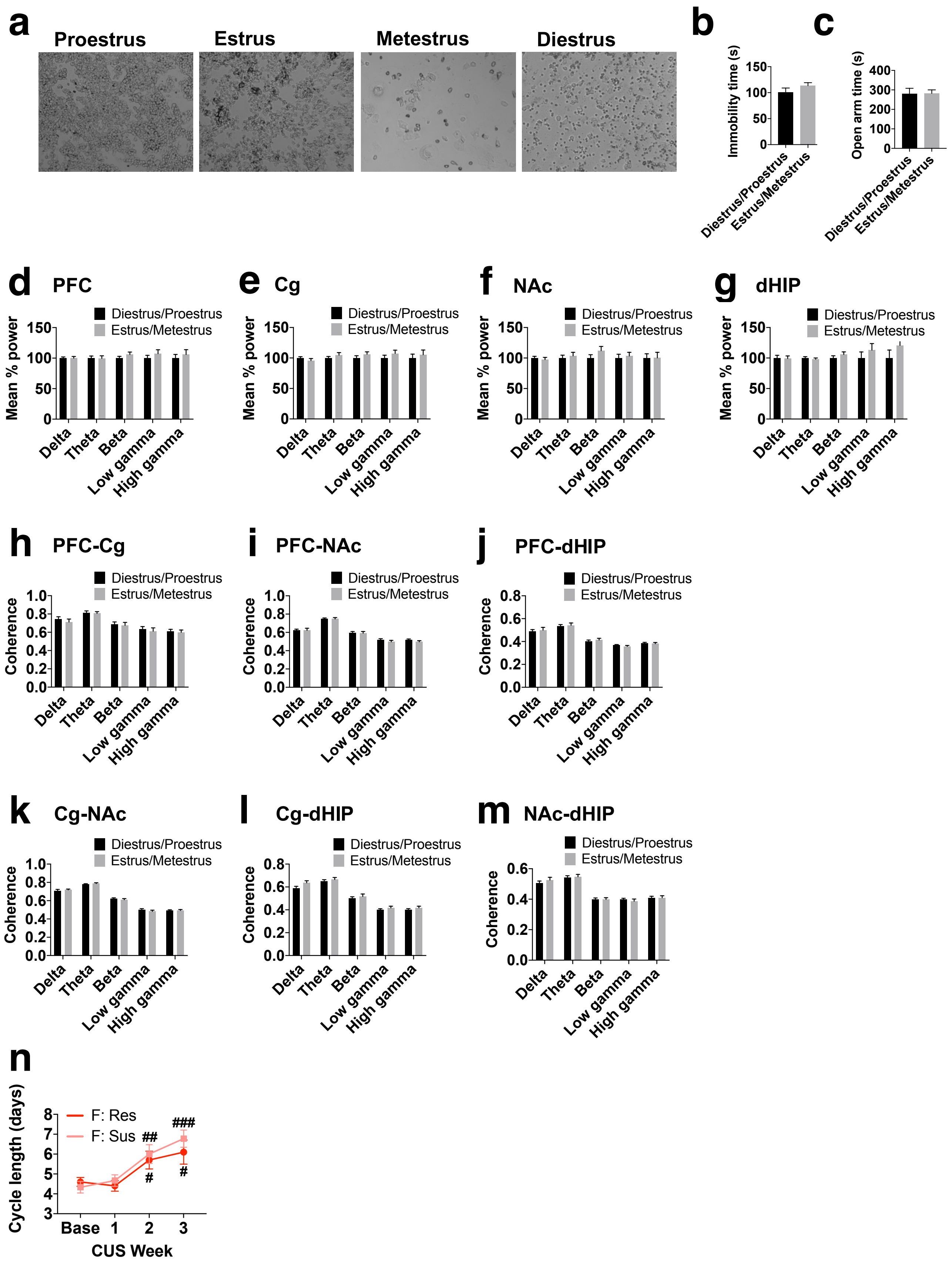

### Supplemental Fig. 4

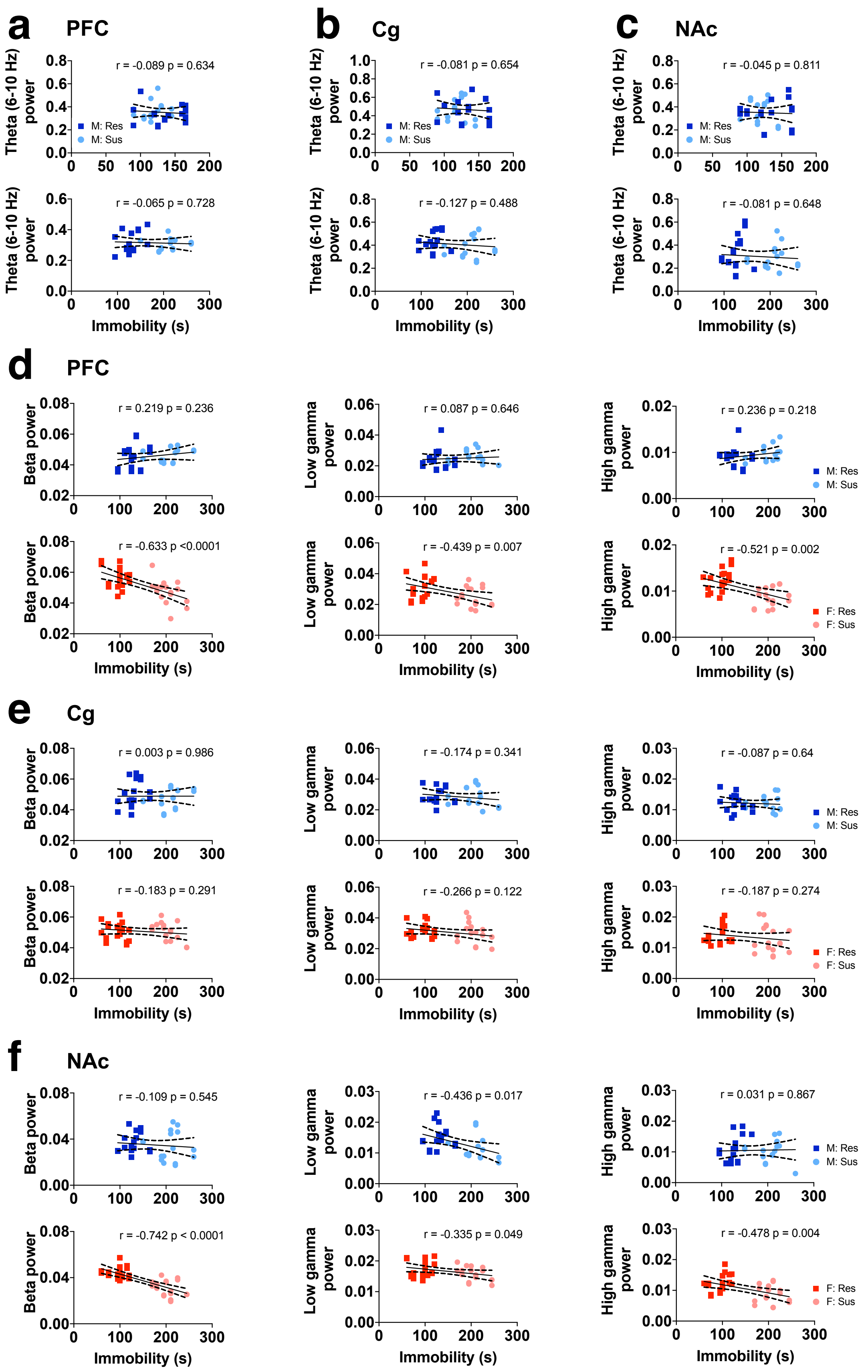

**Supplemental Fig. 5**

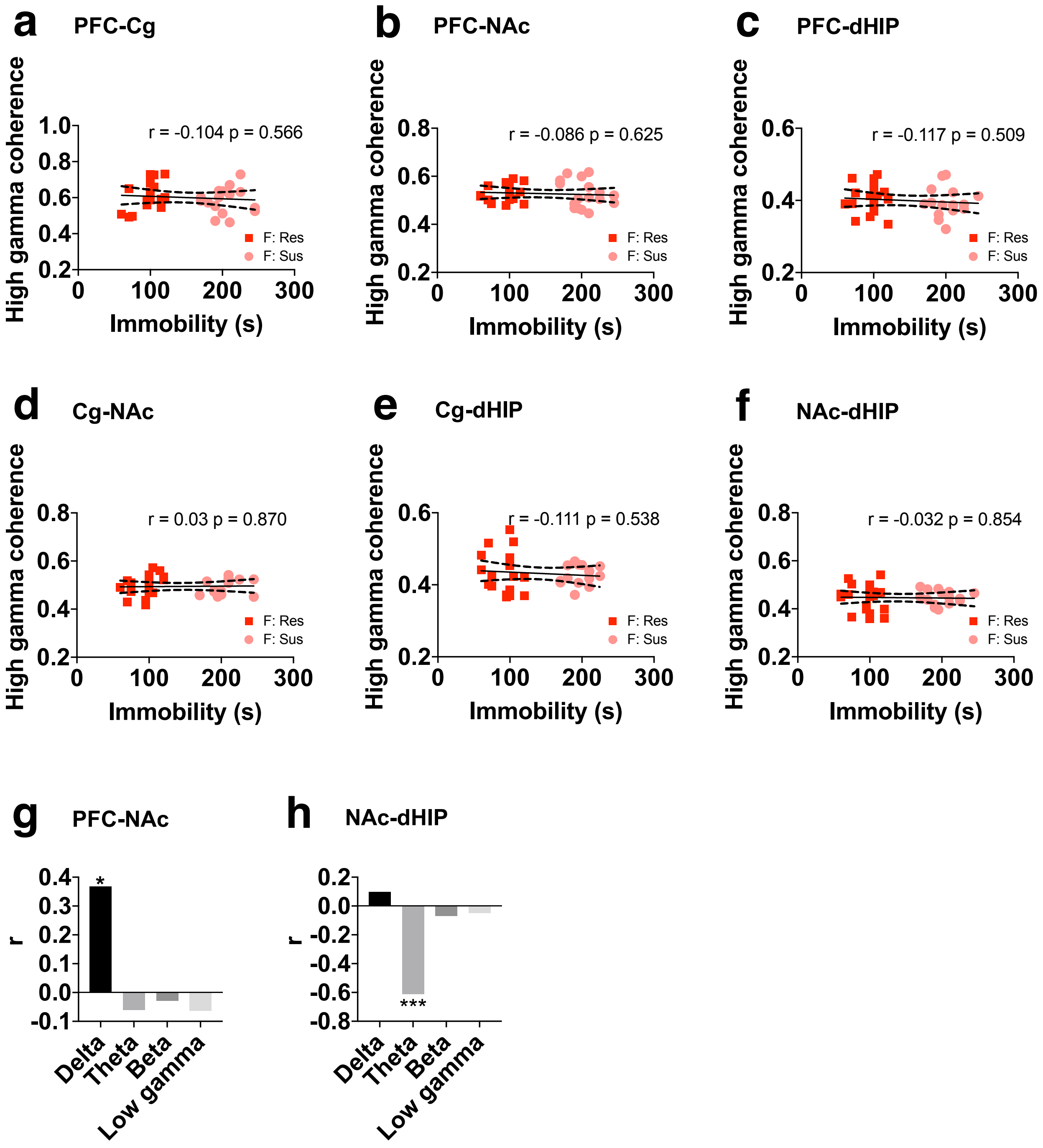

**Supplemental Fig. 6**

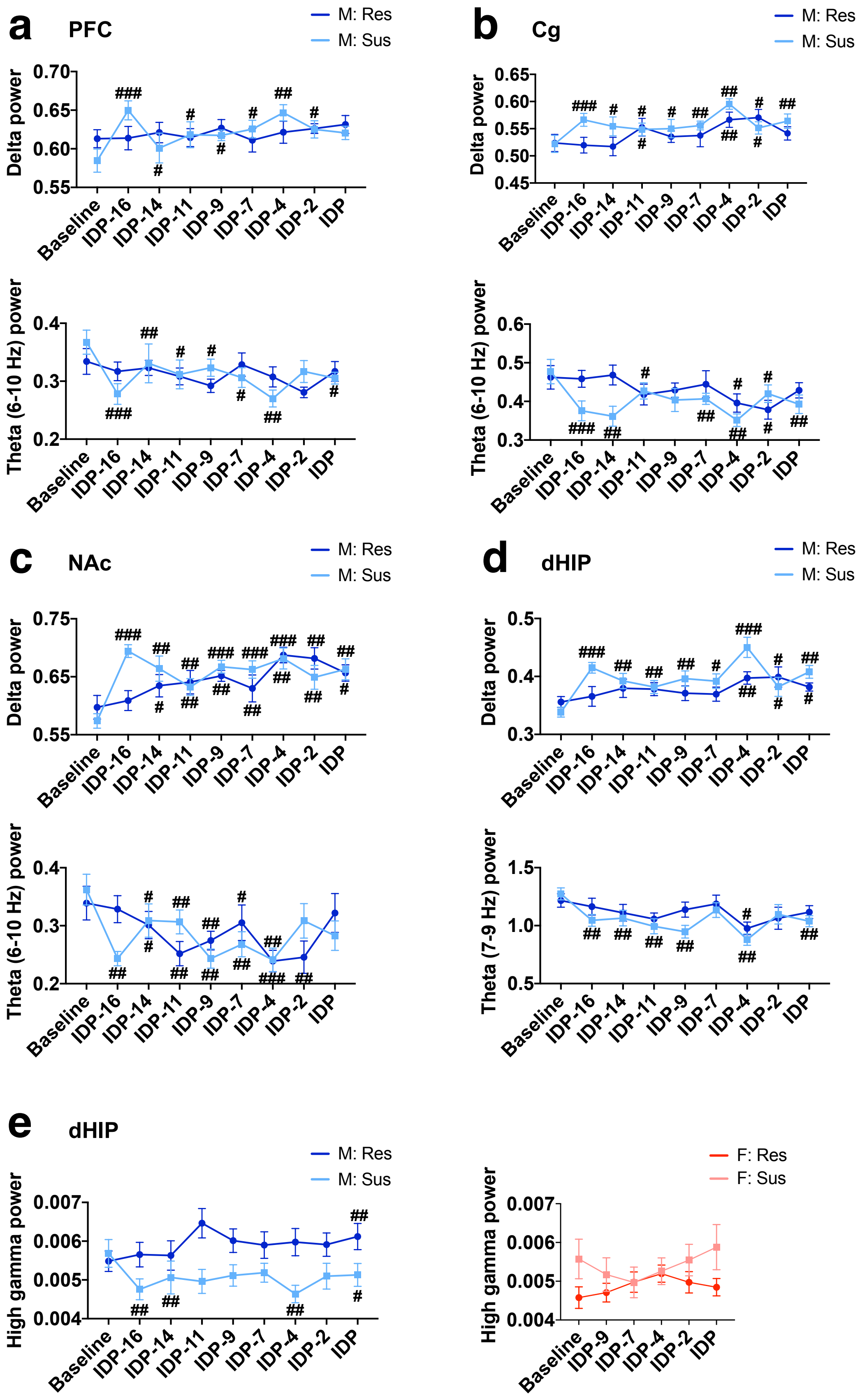

**Supplemental Fig. 7**

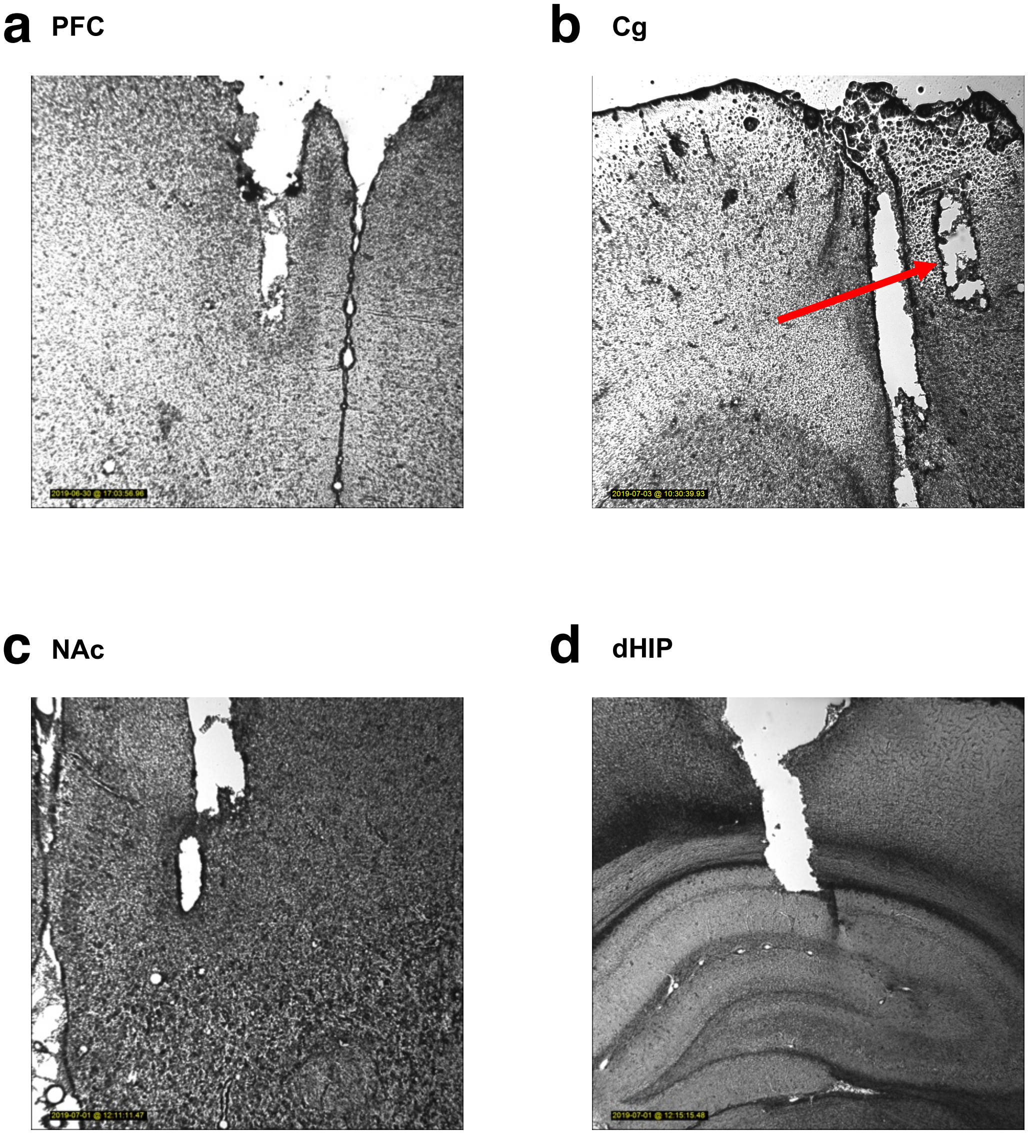
